# Calcium-responsive transactivator (CREST) toxicity is rescued by loss of PBP1/ATXN2 function in a novel yeast proteinopathy model and in transgenic flies

**DOI:** 10.1101/415927

**Authors:** By Sangeun Park, Sei-Kyoung Park, Naruaki Watanabe, Tadafumi Hashimoto, Takeshi Iwatsubo, Tatyana A. Shelkovnikova, Susan W. Liebman

## Abstract

Proteins associated with familial neurodegenerative disease often aggregate in patients’ neurons. Several such proteins, e.g. TDP-43, aggregate and are toxic when expressed in yeast. Deletion of the ATXN2 ortholog, *PBP1,* reduces yeast TDP-43 toxicity, which led to identification of ATXN2 as an amyotrophic lateral sclerosis (ALS) risk factor and therapeutic target. Likewise, new yeast neurodegenerative disease models could facilitate identification of other risk factors and targets. Mutations in *SS18L1,* encoding the calcium-responsive transactivator (CREST) chromatin-remodeling protein, are associated with ALS. We show that CREST is toxic in yeast and forms nuclear and occasionally cytoplasmic foci that stain with Thioflavin-T, a dye indicative of amyloid-like protein. Like the yeast chromatin-remodeling factor SWI1, CREST inhibits silencing of *FLO* genes. Toxicity of CREST is enhanced by the [*PIN*^+^] prion and reduced by deletion of the *HSP104* chaperone required for the propagation of many yeast prions. Likewise, deletion of *PBP1* reduced CREST toxicity and aggregation. In accord with the yeast data, we show that the Drosophila ortholog of human ATXN2, dAtx2, is a potent enhancer of CREST toxicity. Downregulation of dAtx2 in flies overexpressing CREST in retinal ganglion cells was sufficient to largely rescue the severe degenerative phenotype induced by human CREST. Overexpression caused considerable co-localization of CREST and PBP1/ATXN2 in cytoplasmic foci in both yeast and mammalian cells. Thus, co-aggregation of CREST and PBP1/ATXN2 may serve as one of the mechanisms of PBP1/ATXN2-mediated toxicity. These results extend the spectrum of ALS associated proteins whose toxicity is regulated by *PBP1/ATXN2*, suggesting that therapies targeting ATXN2 may be effective for a wide range of neurodegenerative diseases.

**Author summary:** Mutations in the calcium-responsive transactivator (CREST) protein have been shown to cause amyotrophic lateral sclerosis (ALS). Here we show that the human CREST protein expressed in yeast forms largely nuclear aggregates and is toxic. We also show that the HSP104 chaperone required for propagation of yeast prions is likewise required for CREST toxicity. Furthermore deletion of HSP104 affects CREST aggregation. ATXN2, previously shown to modify ALS toxicity caused by mutations in the TDP-43 encoding gene, also modifies toxicity of CREST expressed in either yeast or flies. In addition, deletion of the yeast ATXN2 ortholog reduces CREST aggregation. These results extend the spectrum of ALS associated proteins whose toxicity is regulated by *ATXN2*, suggesting that therapies targeting ATXN2 may be effective for a wide range of neurodegenerative diseases.

## Introduction

Mutations in an increasing number of human genes have been found to cause familial neurodegenerative disease [1, 2]. Proteins encoded by these genes are often soluble in healthy individuals, but form insoluble amyloid-like aggregates that seed further aggregation in the neurons of patients with disease. For example, such conformational changes have been seen for: Aβ, associated with Alzheimer’s disease; α-synuclein with Parkinson’s disease; TDP-43, FUS and others with amyotrophic lateral sclerosis (ALS) and frontotemporal dementia (FTD); and huntingtin with Huntington’s disease. Wild-type and mutant forms of these proteins are respectively associated with aggregates in sporadic and familial forms of the diseases. Overexpression of either wild-type or mutant protein causes toxicity, although the mutant forms are often toxic at lower concentrations than the wild-type [3].

Conformational change of several yeast proteins from a soluble state to insoluble self-seeding aggregates, called prions, causes transmissible phenotypic changes [4–15]. Furthermore, the presence of one prion aggregate enhances the *de novo* appearance of heterologous prions. For example, the endogenous yeast prion [*PIN*^+^] (sometimes called [*RNQ^+^*]), which is an amyloid form of the RNQ1 protein, promotes the *de novo* aggregation of the SUP35 protein to form the [*PSI*^+^] prion. This could occur by cross-seeding or by sequestration of proteins such as chaperones by the amyloid [*PIN*^+^] prion [8,16–27].

Yeast has proved to be useful in the study of disease-specific proteins that form prion-like aggregates [28–35]. When human proteins associated with aggregation in neurodegenerative disease were expressed in yeast, they formed aggregates and caused toxicity. Curiously the toxicity of several of these proteins, e.g. TDP-43, FUS or huntingtin, is enhanced by the presence of the [*PIN*^+^] prion [28,36,37]. These yeast models have allowed the identification of yeast genes that either alter the disease protein’s aggregation or enhance or reduce its toxicity. Remarkably, human homologs of these yeast modifier genes have confirmed, and identified new human risk factors for Alzheimer’s (*PICALM, XPO1*, *ADSSL1* and *RABGEF1*) [38–40], Parkinson’s (*PARK9*) [38, 41]) and ALS (*ATXN2*) [42–46].

Indeed, the discovery in yeast that deletion of the *ATXN2* (Ataxin-2) ortholog, *PBP1,* reduced TDP-43 toxicity, led to the recent exciting finding that reduction in ATXN2 levels dramatically lowers neurotoxicity in ALS mouse models [43]. Thus the establishment of new yeast neurodegenerative disease models may lead to the identification of new risk factors for human disease as well as provide screening platforms for drug discovery.

Recently, mutations in *SS18L1* have been shown to cause ALS [47, 48]. *SS18L1* encodes the calcium-responsive transactivator (CREST) protein which is an essential component of the nBAF neuron-specific chromatin remodeling complex that is related to the yeast SWI/SNF chromatin remodeling complex [49–51]. Overexpression of CREST inhibits neurite outgrowth in cultured neurons and causes retina degeneration in transgenic Drosophila [48, 52]. Also, CREST co-immunoprecipitates with FUS in neuron lysates [48, 52]. In cultured cells, transformed CREST forms predominantly nuclear dots with some cytoplasmic foci. Also, like other ALS associated proteins, CREST is recruited to stress-induced ribonucleoprotein particle (RNP) granules [52, 53]. While many proteins must reversibly aggregate in physiological granules as part of their normal cellular function, this ability to aggregate could lead to the formation of pathological aggregates. Indeed, TDP-43 and FUS have prion-like domains and have been found in cytoplasmic aggregates in the neurons of patients with ALS. While no such aggregates have yet been reported for CREST, it has an unstructured prion-like domain predicted to form amyloid-like aggregates [48]. CREST in cell culture was slightly resistant to Triton-X100 but was solubilized by SDS; in addition it formed nuclear dots in a concentration-dependent manner [52].

Here we establish a new ALS model in yeast. We show that this human chromatin remodeling complex component and transcriptional activator is toxic and forms largely nuclear aggregates that stain with Thioflavin T when expressed in yeast. We also show that CREST toxicity is increased by the endogenous yeast [*PIN*^+^] prion and reduced by deletion of the HSP104 chaperone that is required for propagation of many yeast prions [4]. Deletion of the TDP-43 toxicity modifier, *PBP1,* likewise reduces CREST toxicity and this finding was also confirmed in a CREST transgenic fly model. Both *hsp104Δ* and *pbp1Δ* alter CREST aggregation. Finally, overexpression of *PBP1* causes a large portion of CREST to leave the nucleus and co-localize with PBP1 containing cytoplasmic dots in yeast. Likewise, in mammalian cells, cytoplasmic aggregates of CREST sequester ATXN2. Overall, our results establish PBP1/ATXN2 as a potent modifier of CREST toxicity *in vivo*.

## Results

### Expression of human CREST in yeast is toxic

Growth of yeast transformed with control *GAL-GFP* vectors was compared with yeast expressing fusions of human CREST cDNA and GFP from a *GAL* promoter on three types of vectors: a **<u>Y</u>**east ***<u>C</u>****EN* **<u>p</u>**lasmid (YCp) containing a centromere and expressed as one or a few copies in each cell (Fig 1A); a **<u>Y</u>**east **<u>E</u>**pisomal **<u>p</u>**lasmid (YEp) containing elements of the endogenous yeast 2μ episome and expressed in high copy in some cells but absent in other cells (Fig 1B); or a **<u>Y</u>**east **I**ntegrating **<u>p</u>**lasmid (YIp) that was integrated into the genome (Fig 1C). Growth inhibition is evident but slight when CREST-GFP was expressed from a *CEN* vector (Fig 1A) and more inhibition is seen when episomal (Fig 1B) or integrating vectors (Fig 1C) were used. Expression of the *GAL* controlled genes was turned on with a constitutively expressed fusion of the human estrogen receptor hormone-binding domain, the yeast *GAL4* DNA-binding domain and the VP16 viral transcriptional activator, which was activated by the addition of β-estradiol [54]. The high level of β-estradiol used here (2 μM) in glucose media results in more expression from the *GAL* promoter than that obtained by traditional induction on 2% galactose without β-estradiol [55]. Note the reduced induction of CREST caused by galactose relative to β-estradiol results in reduced toxicity (S1 Fig).

**Fig 1.**
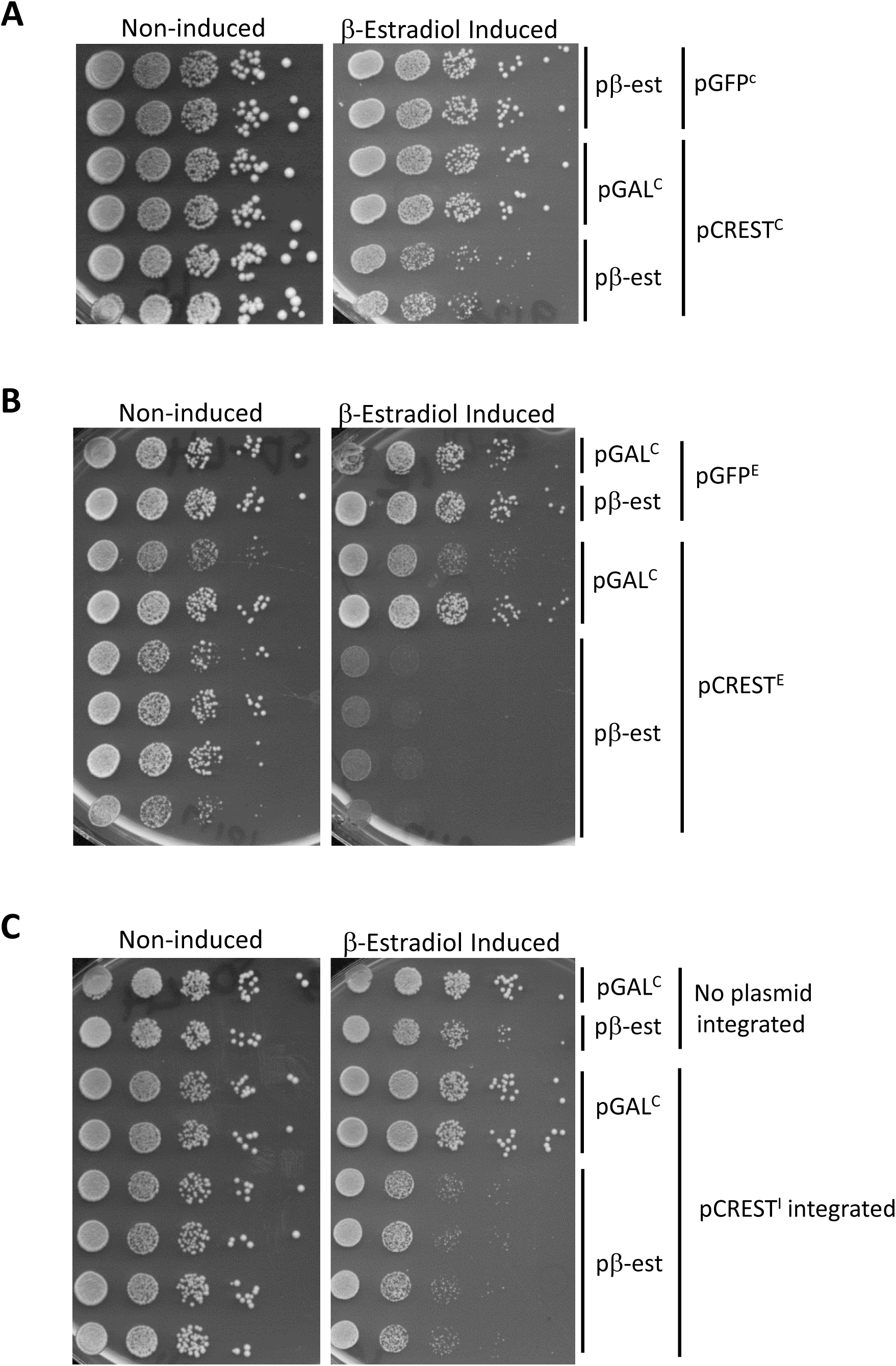
Expression of human CREST in yeast is toxic. Shown is the growth of cells carrying vectors encoding *GAL-CREST-GFP* and a fusion protein that promotes induction of the *GAL* promoter in the presence of estradiol (pβ-est) spotted on plasmid selective glucose plates without estradiol where CREST-GFP expression is *Non-induced*, or with 2 μM β-estradiol where CREST-GFP expression *is β-Estradiol Induced*. Duplicates are independent transformants or integrants. Cells grown on plasmid selective glucose plates were diluted to OD_600_=2 and then serially diluted 10-fold and spotted on the plates shown. The pβ-est plasmid expresses the fusion activator that is responsive to β-estradiol, and YCp*GAL-ccdB* (pGAL^C^) was used as its control. (A) *CREST-GFP expressed from a CEN plasmid inhibits growth.* YCp*GAL-CREST-GFP* (pCREST^C^) and control YCp*-GAL-GFP* (pGFP^C^) vectors were used. (B) *CREST-GFP expressed from a 2μ plasmid inhibits growth.* YEp*GAL-CREST-GFP* (pCREST^E^) and the control YEp*GAL-GFP* (pGFP^E^) vectors were used. (C) *Integrated CREST-GFP is toxic.* Strain with integrated YIp*GAL-CREST-GFP* (pCREST^I^) and parent strain lacking the integrated plasmid but containing control YEp*GAL-GFP* (*LEU2*) was used. Strain 74D-694 [*PIN*^+^] was used for the *CEN* and integrated vector experiments (A and C) while strain BY4741 [*PIN*^+^] was used for the 2μ vector experiments (B).

### CREST forms largely nuclear and occasionally cytoplasmic amyloid-like foci in yeast

Similar to mammalian cells [52], CREST-GFP expressed in yeast largely appeared as dots in nuclei, but smaller cytoplasmic foci were also visible and were more frequent after longer induction times (Fig 2A). Nuclear localization was confirmed by using DAPI stain or a constitutively expressed nuclear HTB1-mCh marker (Figs 2B and S2) which co-localized with CREST-GFP. CREST foci also stained with Thioflavin T that marks amyloid-like protein (Fig 2C). For controls we show TDP-43 and FUS aggregates that, respectively do not and do, stain with Thioflavin T [56]. In addition, the bulk of the CREST protein is found in the pellet of lysates while the non-aggregated proteins, GFP and PGK, remain in the supernatant (Fig 2D).

**Fig 2.**
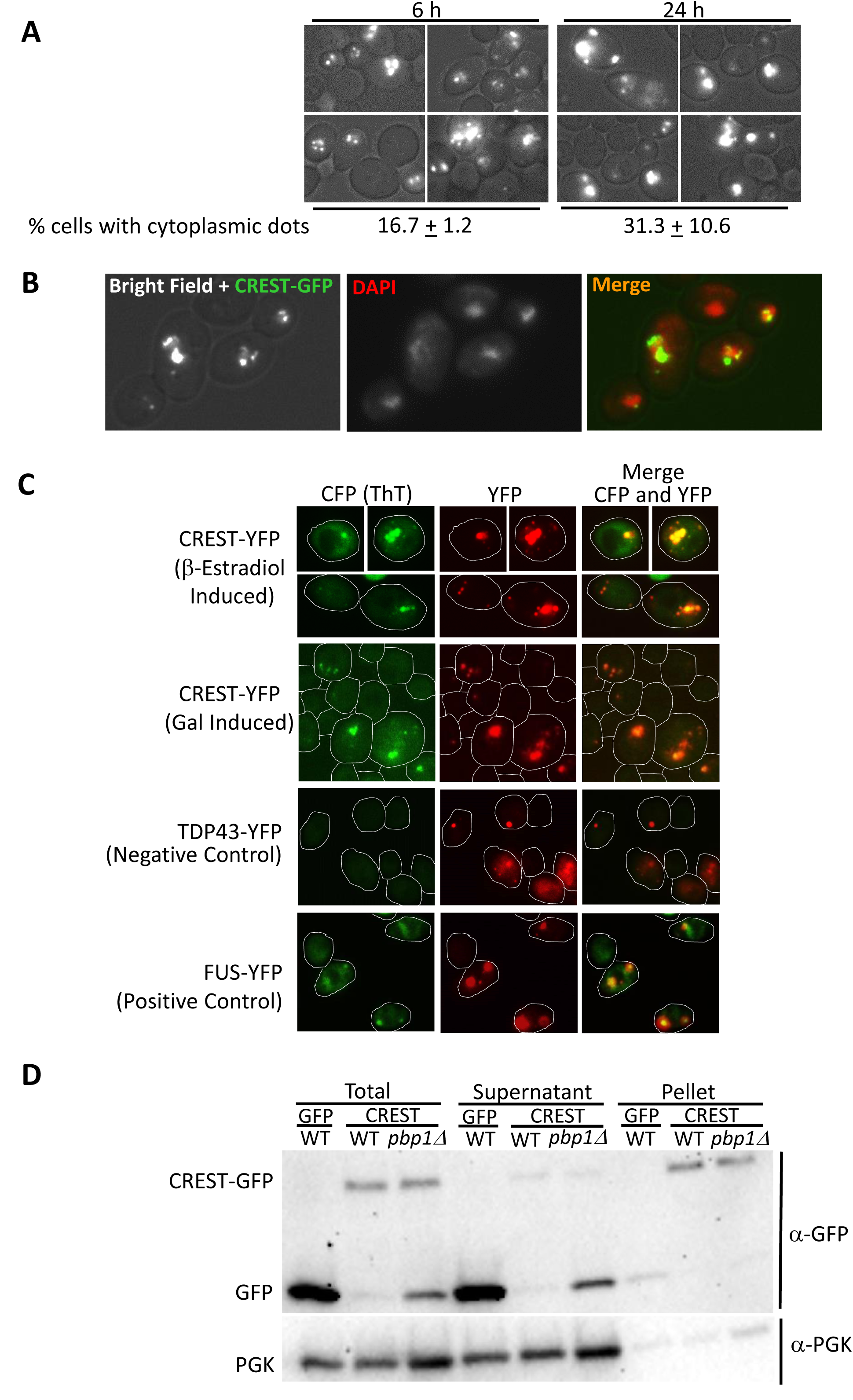
Human CREST expressed in yeast forms largely nuclear dots with occasional cytoplasmic foci. (A) *CREST-GFP expressed in yeast*. BY4741 [*PIN*^+^] transformed with YEp*GAL-CREST-GFP* and pβ-est was induced for 6 or 24 h by growth in plasmid selective glucose media with 2 µM β-estradiol and examined under a fluorescent microscope. Below is shown the % of cells containing CREST-GFP that contained one or more cytoplasmic dots. All of these cells also had nuclear dots. (B) *Co-localization of wild-type CREST-GFP aggregates and DAPI nuclear stain*. Cells in (a) induced for 6 h were stained with DAPI. (C) *CREST aggregates stained with the Thioflavin T (ThT).* Top row: BY4741 transformed YEp*GAL-CREST-YFP* (CREST-YFP) and pβ-est was induced for 6 h with 2 µM β-estradiol. Next three rows strain L3491 transformed with YEp*GAL-CREST-YFP* (CREST-YFP), YCp*GAL-TDP-43-YFP* (TDP-43-YFP) as negative control, or YCp*GAL-FUS-YFP* (FUS-YFP) as a positive control, were grown in plasmid selective SRGAL medium overnight. ThT fluorescence was detected with a CFP filter. CFP was recolored green and YFP recolored red to facilitate visualization of merged pictures. Cell boundaries are drawn. (D) *CREST-GFP is found mostly in pellets in WT but mostly in supernatant of pbp1Δ strains.* Three independent transformants of (*WT*) and (*pbp1Δ*) with pβ-est and either YEp*GAL-CREST-GFP* (CREST) or control YEp*GAL-GFP* (GFP) were grown were grown on plasmid selective glucose medium, induced with 2 µM β-estradiol for 6 h and harvested. Total lysates were separated into *Supernatant* and *Pellet* by centrifugation. Samples were run on SDS-PAGE. Three gels were immunoblotted with anti-GFP, stripped and reprobed with anti-PGK for loading control. All gels showed a ratio of more GFP reactive material in pellets of WT vs. supernatant and the reverse for *pbp1Δ*. Since ratios are used loading differences are not relevant.

When CREST was overexpressed in mammalian cells, it formed large dot-like nuclear structures and was also present in cytoplasmic aggregates [52]. When CREST is expressed in yeast for 5-6 h, most of the protein is found in nuclear dots with just occasional cytoplasmic foci (see Figs 2 and 3 left). However, when cells with nuclear CREST are stressed by incubation at high temperature there is a dramatic increase in the appearance of cytoplasmic foci (Fig 3 middle) that partially co-localize with Ded1-mCh stress granule and EDC3-mCh P-body markers. Following a return to 30°C, cytoplasmic CREST aggregates disappear along with stress granules and P-bodies (Fig 3 right).

**Fig 3.**
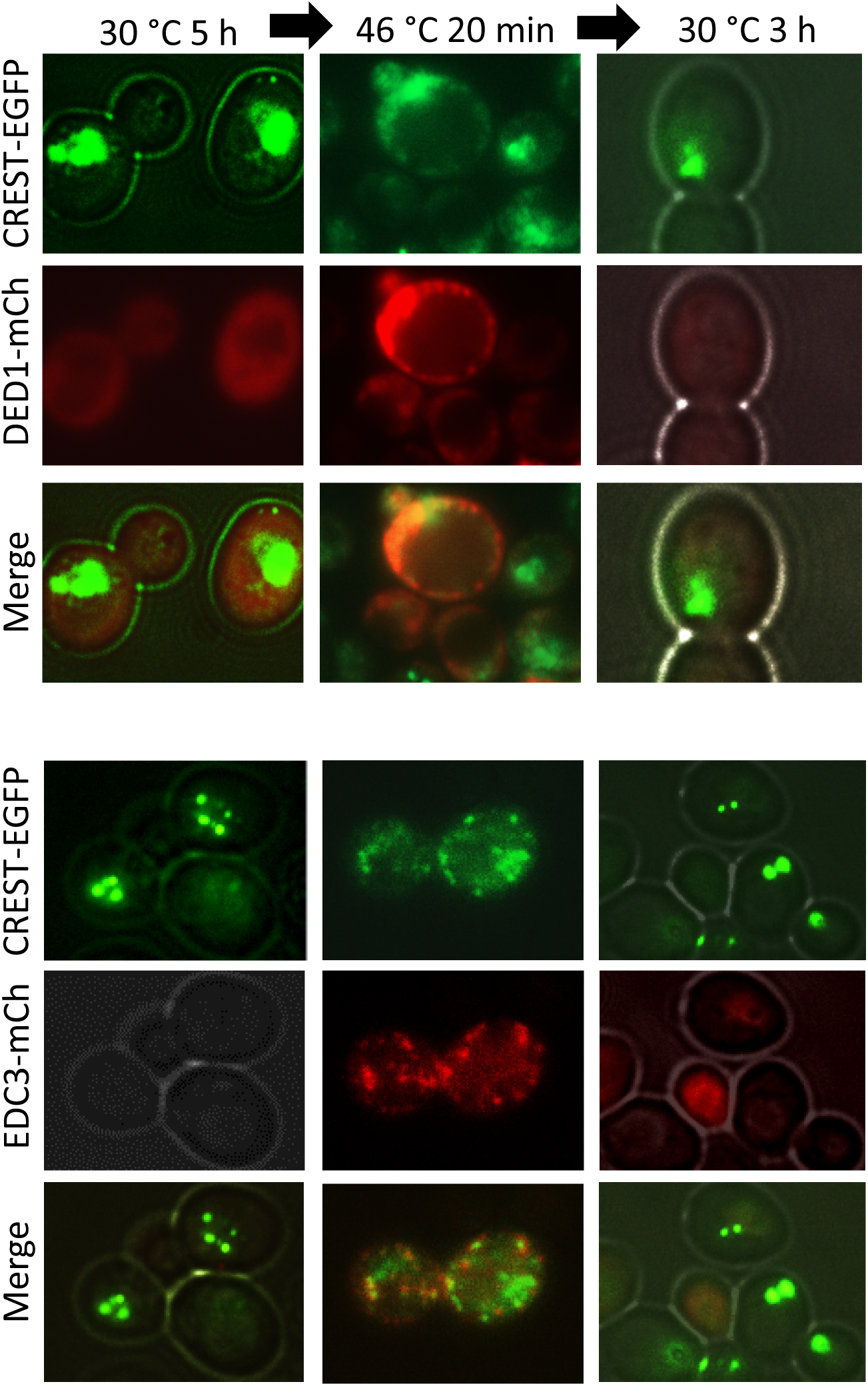
Heat stress causes relocalization of some CREST aggregates from the nucleus to the cytoplasm. CREST partially co-localizes with the DED1 stress granule and EDC3 P-body markers during heat-stress and returns to the nucleus following heat-stress. YCp*DED1-mCh* and YCp*EDC3-mCh* transformants of [*PIN*^+^] 74D-694 with integrated *GAL-CREST-EGFP* were grown in plasmid selective SRaf (2%) to OD600 = 0.3 when CREST-EGFP was induced by addition of 2% galactose for 5 h. Cells were then stressed for 20 min at 46°C. CREST-EGFP, DED-mCh and EDC3-mCh were examined before (30°C), during (46°C) and 3 h following (30°C) heat stress.

### The presence of the [*PIN*^+^] prion or HSP104 enhances toxicity of CREST

Since the presence of the [*PIN*^+^] prion causes enhanced toxicity of the human genes huntingtin/polyQ, TDP-43 and FUS [28,36,37,57], we asked if [*PIN*^+^] would likewise increase toxicity of CREST and found that it did (Fig 4A). However, CREST and [*PIN*^+^] aggregates did not co-localize (Fig 4B). Also, CREST aggregated in both [*PIN*^+^] and [*pin*^-^] cells (Figs 2A and 4D). Interestingly, CREST toxicity was reduced even further when the *HSP104* chaperone required for the propagation of many yeast prions [4, 58] was deleted in [*pin*^-^] cells (Fig 4C left). Deletion of *HSP104* also caused the appearance of many nuclei with diffuse CREST-GFP with or without dots instead of nuclear CREST-GFP dots on a black background, and reduced the number of cells showing any cytoplasmic CREST aggregation (Fig 4D). CREST-GFP was found largely in the pellet in lysates of both *hsp104Δ* and *HSP104* yeast (Fig 4E). The higher level of CREST-GFP in the *hsp104Δ* cultures (Fig 4C right) could reflect the reduced toxicity of CREST.

**Fig 4.**
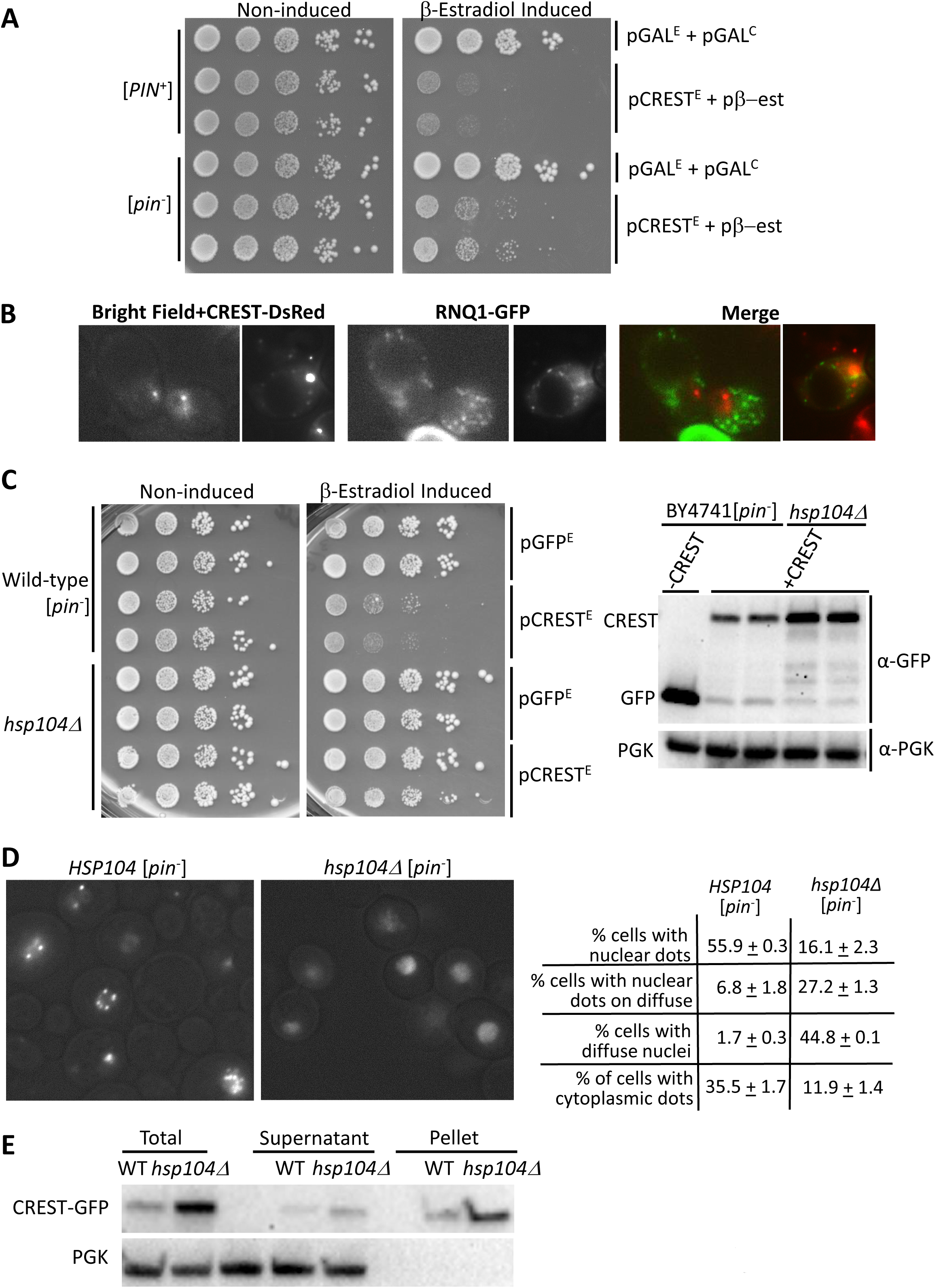
Toxicity of CREST is reduced by the absence of the [*PIN^+^*] prion, and is further reduced by the absence of HSP104. (A)*The [PIN^+^] prion enhances toxicity of CREST in yeast.* Shown is the growth (as in Fig 1) of [*PIN*^+^] and [*pin*^-^] BY4741 cells carrying YEp*GAL-CREST-GFP* (pCREST^E^) or control YEp*GAL-GFP* (pGFP^E^) and the pβ-est plasmid or empty control YCp*GAL-ccdB* (pGAL^C^) on plasmid selective glucose media in the presence (β-Estradiol Induced) and absence (Non-induced) of β-estradiol. (B) *[PIN^+^] and CREST aggregates do not co-localize.* [*PIN*^+^] BY4741 carrying pβ-est, YEp*GAL-CREST-DsRed* (CREST-DsRed) and YCp*RNQ-GFP* (RNQ1-GFP) were grown for 2 d on plates containing 50 μM CuSO_4_ and 2 μM β-estradiol to respectively induce RNQ1-GFP and CREST-DsRed and were then examined with a fluorescence microscope. (C) *The absence of HSP104 reduces CREST toxicity.* Left: Independent transformants of *HSP104* (Wild-type) or *hsp104Δ* versions of [*pin*^-^] BY4741 carrying pβ-est and YEp*GAL-CREST-GFP* (pCREST^E^) or control YEp*GAL-GFP* (pGFP^E^) vectors were serially diluted and spotted on plasmid selective glucose without (Non-induced) or with 2 μM β-estradiol (β-estradiol Induced). Right: Same transformants were grown in liquid plasmid selective dextrose media and induced with β-estradiol for 6 h. Blots probed with GFP antibody to detect CREST were stripped and reprobed with phosoglycerate kinase (PGK) antibody to detect the PGK loading control. (D) *CREST aggregation is reduced in the absence of HSP104. HSP104* (Wild-type) or *hsp104Δ* versions of [*pin*^-^] BY4741 carrying pβ-est and YEp*GAL-CREST-GFP* grown on plates for 3 d in plasmid selective dextrose medium containing 2 µM β-estradiol were examined with a fluorescence microscope. Among cells expressing CREST-GFP we quantified the percent of cells with: nuclear fluorescent dots, nuclear fluorescent dots on a diffuse nuclear background, nuclei with just diffuse fluorescence and one or more cytoplasmic fluorescent dots. (E) *CREST sediments largely in the pellet in lysates of HSP104-WT or hsp104Δ cells.* Transformants grown in liquid plasmid selective dextrose media and induced with β-estradiol for 6 h were harvested. Lysates were separated into *Supernatant* and *Pellet* factions by centrifugation and samples were run on SDS-PAGE. Two gels immunoblotted with anti-GFP, stripped and reprobed with anti-PGK for loading control gave identical results.

### Expression of human CREST activates yeast *FLO* genes and causes flocculation

While working with *GAL-CREST-GFP* and *GAL-GFP* transformants, we noticed that expression of CREST-GFP caused cells in liquid culture to settle very rapidly compared to the GFP controls. In a measured experiment, we found that overexpression of CREST caused flocculation (cell clumping), while overexpression of TDP-43 or FUS did not (Fig 5A).

Since CREST increased yeast flocculence just as the non-prion form of the SWI1 chromatin remodeling protein does [59], we hypothesized that CREST, like SWI1, remodels yeast chromatin structure causing a release of silencing of telomerically located and epigenetically controlled [60] *FLO* genes, resulting in flocculation. To test this we examined the effect of CREST on the telomerically located *FLO1* and the epigenetically controlled *FLO11* promoters, using *ura3Δ* strains with either the *FLO1* or *FLO11* open reading frames (ORFs) replaced with the *URA3* ORF reporter [59]. In the absence of CREST these strains were auxotrophic for uracil indicating that both the *FLO1* and *FLO11* promoters were off. However, upon CREST expression on galactose medium, both strains gained the ability to grow in the absence of uracil, indicating that the *FLO1* and *FLO11* promoters were indeed both activated by CREST (Fig 5B).

**Fig 5.**
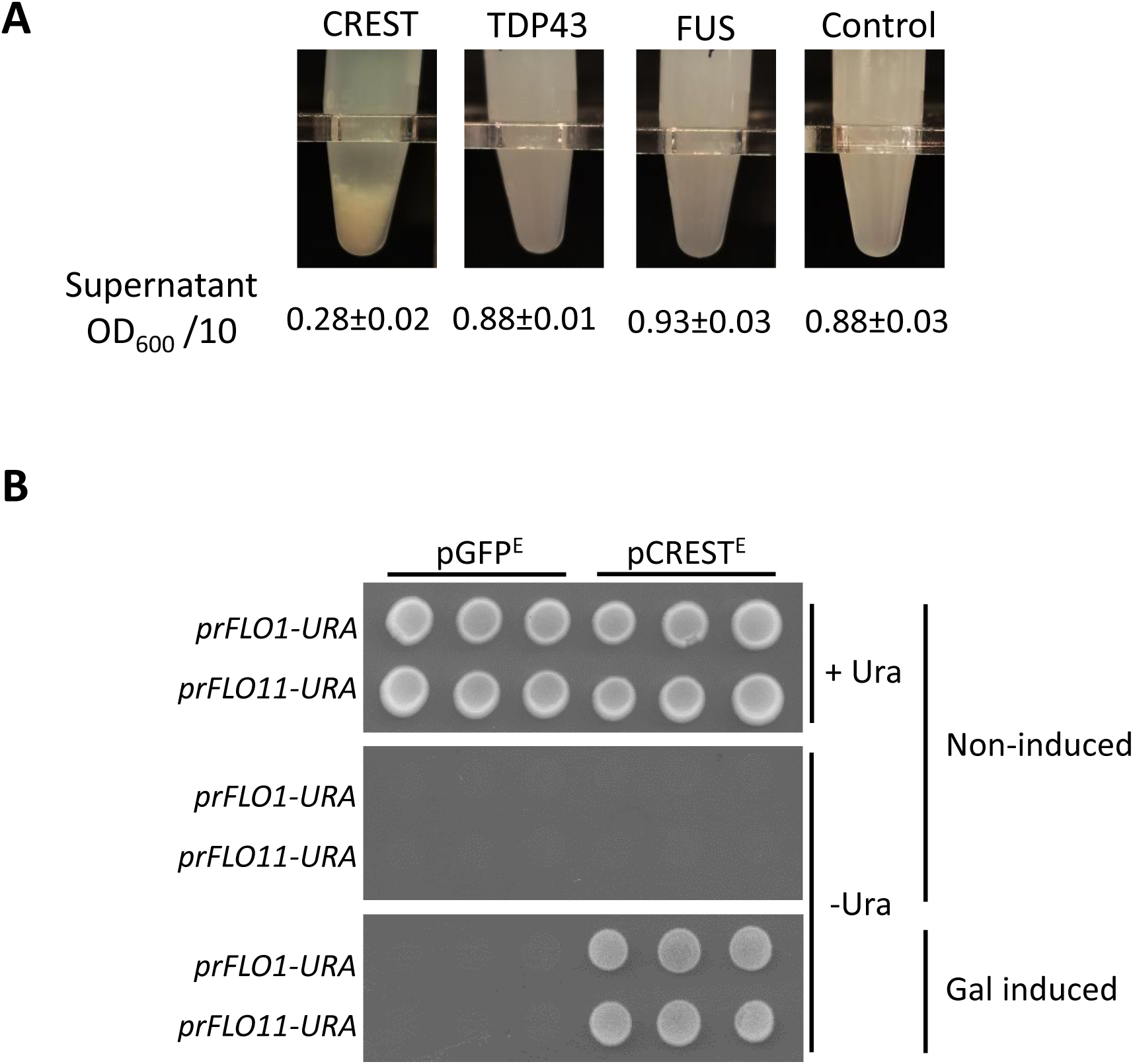
Expression of CREST enhances yeast flocculation. (A) *CREST but not TDP-43 or FUS overexpression enhances flocculation.* Shown are transformants of BY4741 [*PIN*^+^] with: YCp*GAL-CREST-GFP* (CREST), YCp*GAL-TDP-43-YFP* (TDP-43), YCp*GAL-FUS-YFP* (FUS) or control YCp*-GAL-GFP* with the average and standard error of OD_600_ of 10X diluted supernatants after cells of 3 independent transformants were allowed to settle for 15 min. (B) *Expression of CREST activates the FLO1 and FLO11 promoters.* Strains deleted for *URA3* and containing either the *FLO1*(L3580) or *FLO11(*L3582) open reading frames (ORFs) replaced with the *URA3* ORF reporter, respectively *prFLO1:URA and prFLO11:URA* were transformed with YEp*GAL-CREST-GFP* (pCREST^E^) or control YEp*GAL-GFP* (pGFP^E^). Three independent transformants of each were spotted on plasmid selective glucose media (Non-induced), with (+Ura) or without (-Ura) uracil or plasmid selective -Ura galactose media (Gal Induced).

### Unlike [*PIN*^+^] aggregates, CREST dots do not seed SUP35 prion formation

Considerable evidence suggests that amyloid-like aggregates of one protein can facilitate the *de novo* aggregation of certain heterologous proteins. Such a phenomenon could be an important risk factor for disease. While there are no yeast proteins that are homologous to the human CREST, the yeast protein with the most similarity to CREST, using a BLAST search, is RNQ1, the [*PIN*^+^] protein. This is likely due to the extensive QN-rich regions in both proteins. Thus, we asked if aggregates of CREST, like [*PIN*^+^] and other QN-rich aggregates, could facilitate the *de novo* aggregation of SUP35 to form the [*PSI*^+^] prion. In [*PIN*^+^] but not [*pin*^-^] cells, transient overexpression of SUP35NM-GFP causes the appearance of large fluorescent SUP35NM-GFP rings and converts [*psi*^-^] cells into [*PSI*^+^] [8,17,55,61]. Here we detect conversion into [*PSI*^+^], which causes readthrough of nonsense codons, by the appearance of adenine prototrophy despite the presence of the nonsense mutation *ade1-14.* We found that unlike [*PIN*^+^], or overexpression of a variety of QN-rich yeast proteins [8], overexpression of CREST did not cause the appearance of fluorescent SUP35NM-GFP rings or adenine prototrophy. This was true even when *PBP1* was also overexpressed (S3 Fig), which, as described below in the last research section, drives CREST-DsRED aggregates into the cytoplasm.

### PBP1/dAtx2 enhances toxicity of CREST in yeast and flies

Overexpression of yeast’s *PBP1*, a homolog of human *ATXN2*, enhances, while *PBP1* deletion reduces, the toxicity of TDP-43 expressed in yeast [42]. This led to the discovery that ATXN2 intermediate-length polyglutamine expansions, which are associated with increased ATXN2 protein levels, are a risk factor for ALS [42]. Work in a variety of model organisms including mice, now suggests that *ATXN2* is a modifier of some neurodegenerative diseases [43].

We thus tested the effect of *PBP1* level on the toxicity of CREST in yeast. Overexpression of *PBP1* itself was toxic in our system (Fig 6A) so the effect on CREST toxicity could not be determined. However, we clearly show that deletion of *PBP1* reduces toxicity in either a [*PIN*^+^] or [*pin*^-^] background (Fig 6B) and this is associated with an increase in CREST-GFP solubility (Fig 2D). Indeed, lysates of *pbp1Δ* cells expressing CREST-GFP had more GFP reactive protein in the supernatant vs. pellet fractions while the reverse was true for *PBP1* lysates. Furthermore the CREST-GFP in *pbp1Δ* supernatant showed a lot of degradation compared with CREST-GFP in the pellet or in either supernatant or pellet fractions of the isogenic *PBP1* strain (Fig 2D). This is likely because soluble CREST-GFP is more susceptible to degradation than aggregated CREST-GFP found in the pellet. We also saw a significant reduction in the number of cells with cytoplasmic CREST aggregates in the presence of *pbp1Δ* (Fig 6C). Taken together the data imply that a smaller fraction of total CREST-GFP is aggregated in either nuclei or cytoplasm the absence of PBP1.

**Fig 6.**
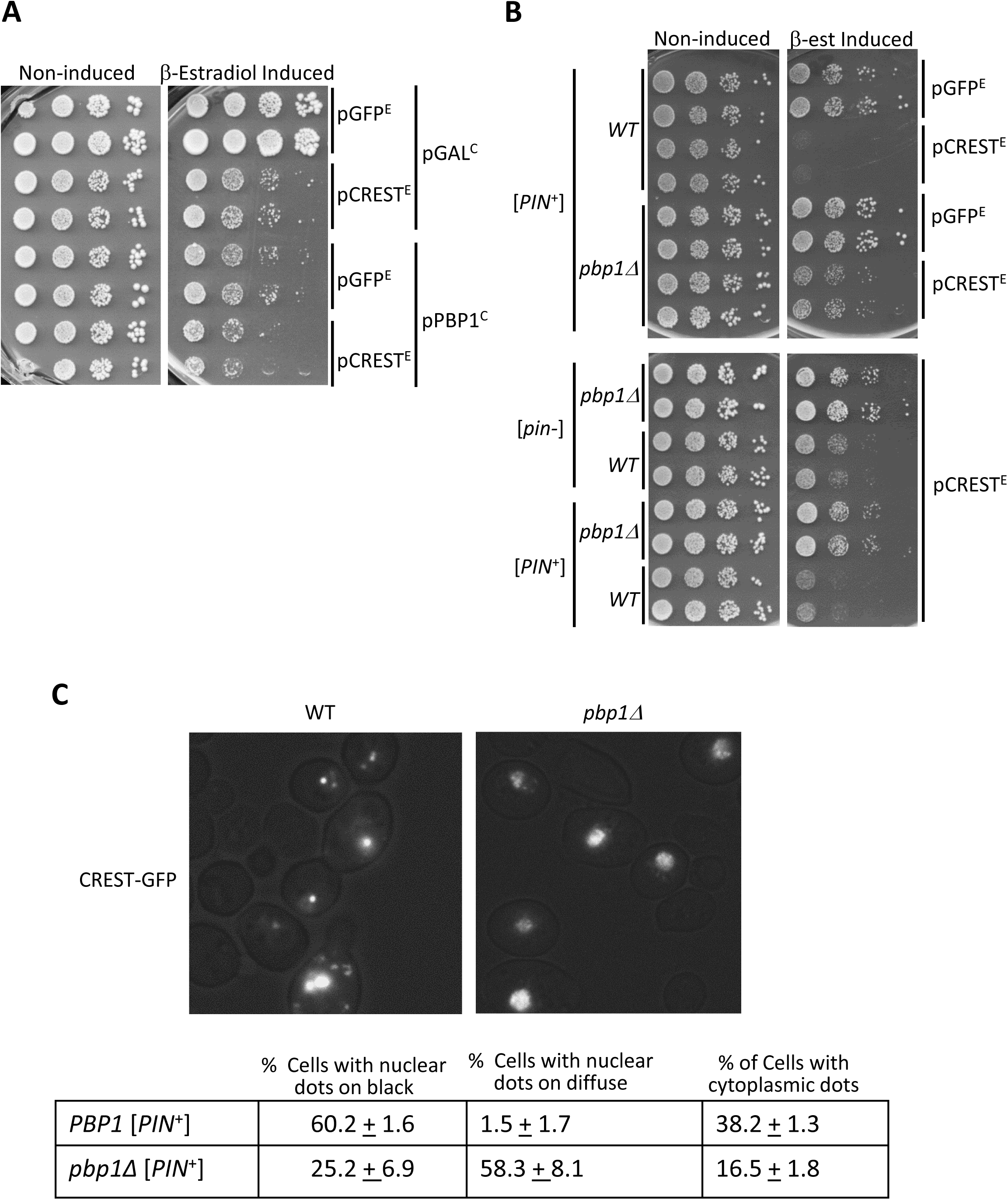
Effects of *PBP1* on CREST. (A) *Overexpression of PBP1 is toxic.* Serial dilutions of [*PIN*^+^] carrying pβ-est and YEp*GAL-CREST-* GFP (pCREST^E^) or control YEp*GAL-GFP* (pGFP^E^) and YCp*GAL-PBP1-GFP* (pPBP1^C^) or control YCp*GAL* (pGAL^C^), were spotted on plasmid selective dextrose (Non-inducing) or β-estradiol containing (β-estradiol Induced) plates and photographed after 3 d at 30°C. (B) *Deletion of PBP1 reduces CREST toxicity.* Independent transformants of [*PIN*^+^] or [*pin^-^*] without (*WT*) and with the deletion (*pbp1Δ*) carrying YEp*GAL-CREST-GFP* (pCREST^E^) or empty GFP control YEp*GAL-GFP* (pGFP^E^) and all carrying pβ-est expressing the fusion activator responsive to β-estradiol, were spotted and grown on plasmid selective media with (β-est Induced), and without (Non-induced), β-estradiol. (C) *Deletion of PBP1 alters CREST aggregation.* Transformants were grown for 2 d on medium with β-estradiol and were then examined. Counts were performed on 3 independent transformants each to determine the fraction of cells with CREST-GFP that had nuclei with dots on a black background, nuclei with dots on a diffuse background and one or more cytoplasmic dots.

In order to validate the observed effect of the ATXN2 ortholog PBP1 in an independent *in vivo* model, we used transgenic *Drosophila* which possesses an ortholog of the human *ATXN2* gene, dAtx2. Overexpression of untagged wild-type CREST using GAL4-UAS system in transgenic lines results in severe retinal degeneration [52]. For our studies we used dAtx2 deletion fly line (dAtx2 06490) [62]. Homozygous deletion of dAtx2 was reported to cause disorganized and decreased photoreceptor neurons, but there was no effect in the heterozygous state [62]. We crossed CREST overexpressing (transgenic, tg) flies with heterozygous dAtx2 flies (dAtx2^06490-/+^) and analyzed the double tg progeny. Histological analysis confirmed normal retinal morphology in of dAtx2^06490-/+^ flies, whereas complete disruption of the regularly ordered arrays of photoreceptor neurons was observed in the retina in CRESTwt tg flies, consistent with the previous report [52]. Strikingly, this severe retinal degenerative phenotype in CRESTwt tg flies was largely rescued by downregulation of dAtx2 (CRESTwt tg; dAtx2^06490-/+^ line) (Fig 7A and B). This rescue was not caused by reducing the expression of CREST (Fig 7C). Thus, human CREST toxicity in flies is largely mediated by endogenous dAtx2.

**Fig 7.**
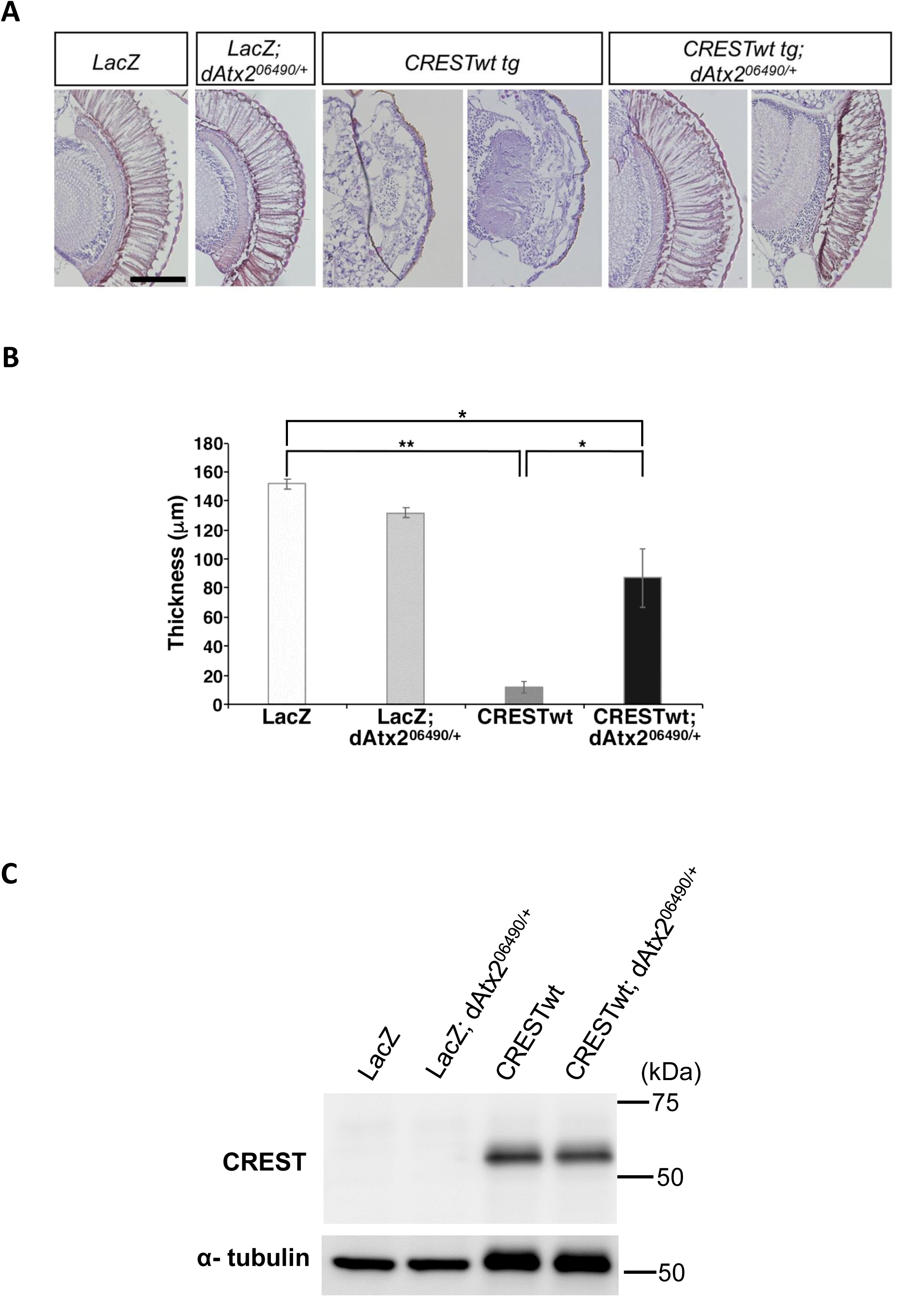
Heterozygous deletion of the Drosophila ortholog of ATXN2, dAtx2, reduced retinal degeneration induced by human CREST. (A,B) *Downregulation of dAtx2 rescues retinal degenerative phenotype in CREST transgenic flies.* H&E staining of retinal sections of 5-day-old flies reveals severe disruption of regularly ordered arrays of photoreceptor neurons in human CRESTwt tg flies, but this phenotype is rescued upon heterozygous deletion of dAtx2 (CRESTwt tg x dAtx2^06490/+^ line). Representative images (A) and quantification (B) for each transgenic fly line are shown. N=10 for LacZ, LacZ; LacZ dAtx206490/+ and CRESTwt tg lines and N=8 for the CRESTwt tg dAtx206490/+ line (3 sections per fly). (**) p<0.01, (*) p<0.05, one-way ANOVA, post-hoc Tukey’s test. Scale bar is 100 um. (C) *Heterozygous deletion of dAtx2 does not alter retinal expression of CREST.* Protein isolated from heads of 5-day-old flies were analyzed as described in materials and methods. α-tubulin was used as control.

### PBP1/ATXN2 and CREST co-localize in cytoplasmic foci in yeast and human cells

Finally, we asked whether potentiated toxicity of CREST in the presence of PBP1 might be linked to the aggregation-prone nature of the two proteins. We found a dramatic increase in cytoplasmic CREST dots when PBP1 was overexpressed in yeast (Fig 8A). This was not due to an increase in CREST protein level (Fig 8B). We also found the CREST-DsRed dots co-localized with PBP1-GFP in cytoplasmic foci (Fig 8C). However these foci do not appear to be RNP granules because they generally lack DED1 and EDC3 (S4 Fig).

**Fig 8.**
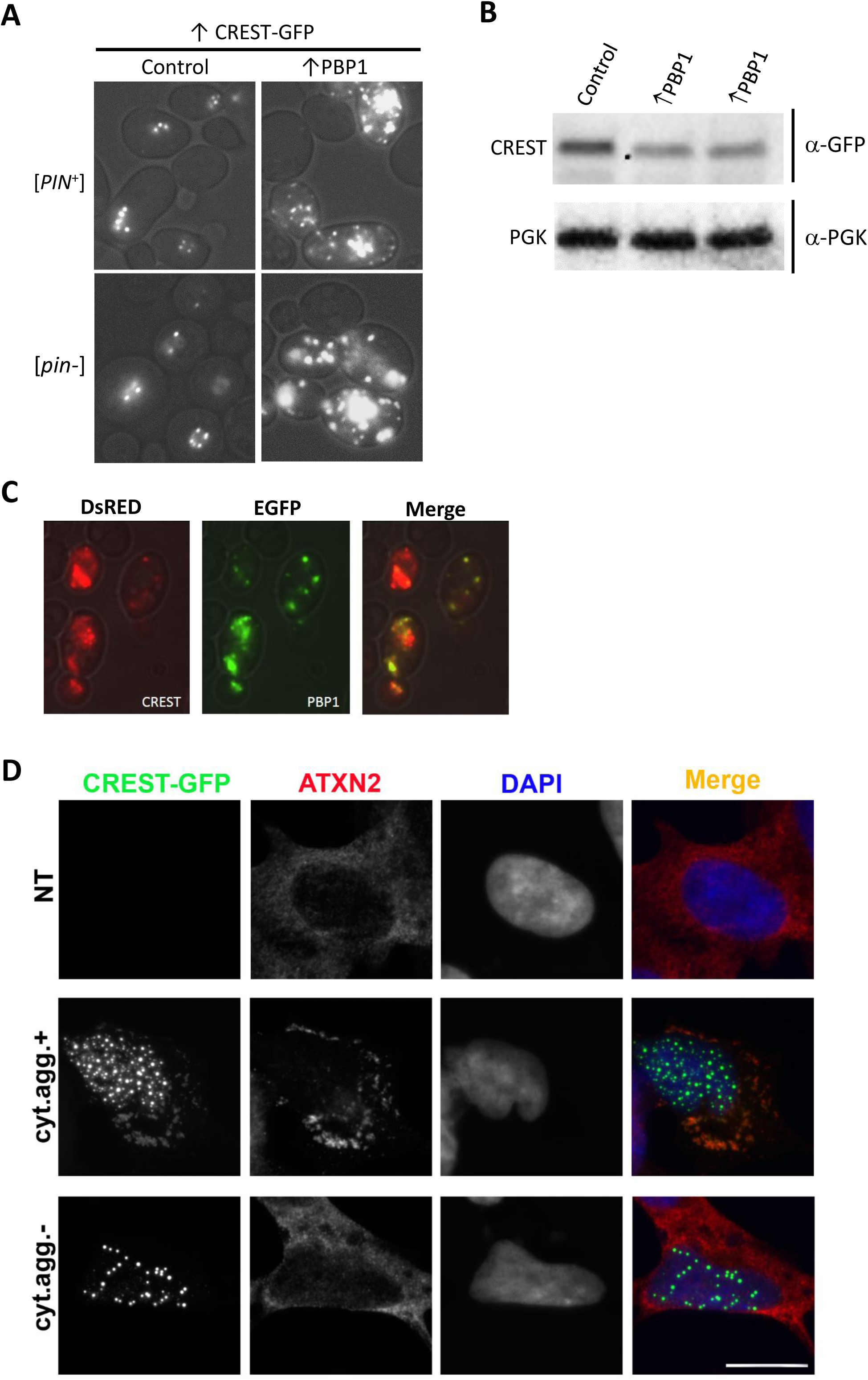
CREST and PBP1/ATXN2 co-localize in cytoplasmic aggregates in yeast and human cells. (A) *Overexpression of PBP1 enhances CREST cytoplasmic aggregation.* Nine co-transformants of [*PIN*^+^] or [pin-] BY4741 with pβ-est and YEp*GAL*-CREST-GFP and either YCp*GAL*-PBP1 (↑PBP1) or YCp*GAL* (Control) were patched and grown on plasmid selective glucose media, velveteen replica-plated and grown on selective plates with β-estradiol for 3 d and examined with a fluorescent microscope. All transformants with ↑PBP1 had a dramatic increase in cytoplasmic foci compared with controls. Shown are representative cells of each type. (B) *PBP1 expression does not increase CREST levels.* Western blot of [*PIN*^+^] transformed with pβ-est and YEp*GAL-CREST-GFP* and either YCp*GAL-PBP1* (↑PBP1) or control vector YCp*GAL*, grown in liquid plasmid selective dextrose media and induced with β-estradiol for 6 hrs. Blots probed with GFP antibody to detect CREST were stripped and reprobed with phosoglycerate kinase (PGK) antibody to detect the PGK loading control. (C). *CREST co-localizes with PBP1 in cytoplasmic foci induced by PBP1 overexpression.* YEp*GAL*-*CREST-DsRED* (CREST) and YCp*GAL*-*PBP1-GFP* (PBP1) were co-transformed into BY4741 [*PIN*^+^]. Transformants patched on plasmid selective glucose medium were velveteen replica-plated, grown on selective SGal for 3 d and examined with a fluorescent microscope. (D) *CREST-GFP overexpressed in neuroblastoma cells sequesters ATXN2 into spontaneously formed cytoplasmic aggregates.* Cells with cytoplasmic CREST aggregates (cyt.agg.+) sequester ATXN2. In cells with only nuclear CREST aggregates (cty.agg.-) or in non-transfected (NT) cells, ATXN2 remains diffuse. Cells were transfected to express CREST-GFP and were analyzed 24 h post-transfection. Scale bar, 10 µM.

We next examined the localization of human ATXN2 in CREST-overexpressing human cells. CREST-GFP forms nuclear aggregates in cells expressing high levels of the protein, and in the ∼10% of cells with the highest levels of CREST-GFP, cytoplasmic aggregates also develop [52]. We found that cytoplasmic CREST-GFP aggregates sequester endogenous ATXN2 in human neuroblastoma cells under these conditions (Fig 8D middle), whereas in non-transfected (NT) and in cells lacking these aggregates, ATXN2 remains diffusely distributed (Fig 8D, top and bottom). Thus, CREST and BP1/ATXN2 co-localize in the cytoplasm, which may serve as one of the mechanisms of PBP1/ATXN2-mediated toxicity.

## Discussion

Our data show that human CREST expressed in yeast shares properties with other ALS-associated proteins and yeast prions: CREST, FUS and yeast prions form amyloid-like foci; CREST, TDP-43 and FUS toxicity and yeast prion appearance are enhanced by the yeast prion [*PIN*^+^]; TDP-43 and CREST toxicity is reduced by deletion of *PBP1*; CREST, TDP-43 and FUS are associated with cytoplasmic RNP granules. Like maintenance of yeast prions, but unlike effects on FUS or TDP-43 [31, 32], CREST toxicity and aggregation is dependent on the presence of HSP104 chaperone.

In mammals, endogenous CREST is a member of a chromatin remodeling complex and thereby activates transcription. When expressed in yeast, we found that CREST releases telomeric silencing and causes flocculation. Thus we propose that even in yeast the heterologous CREST protein remodels chromatin causing transcriptional activation of telomerically located genes including those that cause flocculation (e.g. *FLO1, FLO5, FLO9* and *FLO10*) as well as the epigenetically controlled gene *FLO11*. Likewise, a yeast chromatin remodeling complex protein, SWI1, is required for the expression of telomerically located and epigenetically controlled *FLO* genes [63]. Interestingly, SWI1 can form infectious prion aggregates [12] reminiscent of the amyloid CREST aggregates seen here.

We first described [*PIN*^+^], the prion form of the RNQ1 protein, because its presence allowed the efficient conversion of the SUP35 protein to its prion form, [*PSI*^+^] [8,17,64]. It was later shown that [*PIN*^+^] is also required for the efficient aggregation and toxicity of polyQ in yeast [28]. The mechanisms causing these effects are still unknown. Some data support the hypothesis that the [*PIN*^+^] aggregates of RNQ1 cross-seed aggregation of heterologous proteins such as SUP35 and polyQ. Likewise some, but not all, evidence suggests that the [*PIN*^+^] aggregates bind proteins such as chaperones that would otherwise inhibit aggregation of the heterologous SUP35 or polyQ protein, thereby enhancing their aggregation [18-25,27]

We recently showed that toxicity of TDP-43 and FUS is slightly enhanced by the presence of [*PIN*^+^] [37, 57]. Both of these proteins contain Q/N-rich regions and co-aggregate with polyQ disease protein alleles of huntingtin [65, 66]. Surprisingly, we did not detect a clear effect of [*PIN*^+^] on TDP-43 or FUS aggregation. Likewise, CREST is a Q-rich protein. We show here that the toxicity of CREST is also slightly enhanced by [*PIN*^+^], again with no clear effect on aggregation. Apparently the toxicity of the aggregates can be altered without a noticeable change in their appearance. On the other hand, deletion of either *HSP104* or *PBP1,* which both cause dramatic rescues of CREST toxicity, were each accompanied by reduced frequency of CREST cytoplasmic foci, more diffuse CREST in nuclei and, for *pbp1Δ*, reduced CREST sedimentation.

Surprisingly, despite the fact that we clearly detect more diffuse CREST in the presence of *hsp104Δ*, there was no increase in CREST degradation or in the level of CREST in supernatant fractions as was seen for *pbp1Δ*. Also, immunoblots of *pbp1Δ* lysates reproducibly show that CREST-GFP in the supernatant is degraded much more in the *pbp1Δ* vs. WT *PBP1* strain. CREST-GFP in the pellet fractions was stable in either *PBP1* or *pbp1Δ* strains. This suggests that *pbp1Δ* causes more CREST to remain unaggregated and subject to degradation. Indeed there was more CREST-GFP plus GFP breakdown product in the supernatant vs. pellet in the *pbp1Δ* while the reverse was true for the *PBP1* strain. While *pbp1Δ* also rescues toxicity of TDP-43, effects on TDP-43 aggregation have not been reported [42].

Since the HSP104 chaperone is required to shear the [*PSI*^+^] prion aggregates thereby creating ends where growth occurs, inhibition of HSP104 initially causes [*PSI*^+^] prion polymers to enlarge and, for newly made SUP35-GFP, to not efficiently join the aggregates remaining diffuse instead [67]. Possibly something similar is happening to CREST-GFP in the absence of HSP104. It is also possible that HSP104 is directly involved in CREST aggregate formation.

A recent finding that deletion of *PBP1* causes a reduction in the level of RNQ1 protein [68] could have explained the reduced CREST, TDP-43 and FUS toxicity seen in *pbp1Δ* cells if the lowered RNQ1 level reduced the potency of [*PIN*^+^]. However, we show here that this is not the case because *pbp1Δ* reduces CREST toxicity even in the absence of [*PIN*^+^] (Fig 6E). It remains possible that toxicity is reduced by *pbp1Δ* by reducing RNQ1 levels regardless of [*PIN*^+^] status.

We have shown that the presence of certain QN-rich aggregates can substitute for [*PIN*^+^] and allow the efficient induction of [*PSI*^+^] by overexpression of SUP35 [8]. This fact, plus the observation that RNQ1, the [*PIN*^+^] prion protein, although not a homolog, is the yeast protein most similar to CREST, prompted us to ask if overexpression of CREST could substitute for [*PIN*^+^]. However, CREST did not provide any [*PIN*^+^] activity even when PBP1 was overexpressed driving CREST into the cytoplasm.

Prion aggregates enhance *de novo* formation of heterologous prion, presumably by a cross-seeding mechanism. Curiously, however, if heterologous prion is already present, prion aggregates instead promote loss of the heterologous prion [20]. One mechanism for this has been shown to be prion aggregate binding to, and therefore titrating, chaperones needed to propagate the heterologous prion [24]. Likewise, CREST has been shown to suppress polyQ-mediated huntingtin aggregation and toxicity [69].

Many of the ALS associated proteins, e.g. TDP-43, FUS and TAF15 are soluble nuclear RNA/DNA binding proteins that regulate mRNA splicing and/or stability and contain prion-like unstructured domains and nuclear/cytoplasmic trafficking signals. In neurons of patients with ALS/FTD these proteins have been found in cytoplasmic aggregates instead of their normal nuclear location. This suggests that impaired nucleocytoplasmic transport may be a general mechanism for neurotoxicity [3,70–75]. Indeed, impairing nuclear import of TDP-43 enhances neurotoxicity [76].

While TDP-43 and FUS quickly form cytoplasmic aggregates in yeast with little or no nuclear localization [31,32,37,57,77–79], we found here that CREST appeared in nuclear aggregates with only occasional cytoplasmic foci. While the ALS-associated proteins TDP-43, FUS and SOD1 form aggregates in patients, model systems and *in vitro*, these aggregates are sometimes but not always classical amyloids. In yeast, aggregates of FUS, but not TDP-43, stain with Thioflavin T that is indicative of amyloid-like protein [56]. Although both nuclear and cytoplasmic CREST foci appear to stain with Thioflavin T, it is nonetheless tempting to speculate, by analogy with FUS and TDP-43, that the CREST cytoplasmic foci are more toxic than nuclear CREST. Indeed reduction of CREST-GFP toxicity caused by deletion of either PBP1 or HSP104 also reduced the frequency of cells with cytoplasmic CREST-GFP aggregates.

We found that heat stress induced CREST to leave the nucleus and partially co-localize with RNP granules. This is consistent with previous findings in mammalian cells that CREST becomes mislocalized to the cytoplasm under stressful conditions (oxidative stress) and becomes recruited to cytoplasmic stress granules [52]. Furthermore, as seen for classic stress granule proteins, CREST returned to the nucleus in yeast following the heat stress. Many proteins associated with neurodegenerative disease contain intrinsically disordered regions that are required for their appearance in RNP granules. Evidence has been accumulating that RNP granules are incubators for the formation of pathogenic protein aggregates associated with neurodegenerative disease. It has been proposed that localization of proteins such as TDP-43, FUS and C9ORF72-encoded dipeptide repeat GR50 in high concentration in stress granules likely in a gel form, promotes their conversion into pathogenic amyloid-like aggregates [80–84].

In support of this model, RNP-granule formation has been shown to be strongly associated with neurodegeneration. This was accomplished with the aid of ATXN2, a nucleic-acid-binding protein that is required for RNP-granule formation. Three ALS associated proteins, TDP-43, FUS and C9ORF72-encoded GR dipeptide, have been shown to co-localize with ATXN2 in RNP granules [42, 80] in yeast, human or Drosophila S2 cells. Also, either reducing the level of ATXN2 or deleting one of its intrinsically disordered regions both reduced RNP-granule formation and reduced neurodegeneration caused by TDP-43, C9ORF72 dipeptides or FUS [80,85–87]. These results help explain why expansions of the polyQ regions in ATXN2 that increase the stability, and thus likely the level of ATXN2, are associated with neurodegenerative disease (susceptibility to ALS and type 2 spinocerebellar ataxia).

Here, we report similar effects of PBP1/ATXN2 on CREST: deletion of *PBP1* reduces toxicity of CREST in yeast and fly models of CREST proteinopathy; ATXN2 is recruited into cytoplasmic CREST aggregates in mammalian cells and overexpression of PBP1 causes CREST to co-localize with PBP1 in cytoplasmic foci in yeast. Overexpression of PBP1 has previously been shown to induce the formation of cytoplasmic PBP1 foci [88] that recruit TORC1 thereby down regulating TORC1 function [89]. PBP1 foci may similarly attract CREST which may promote CREST’s conversion to a more toxic form. In sum, we showed that CREST is yet another protein, in addition to TDP-43, that is highly sensitive to ATXN2 dosage indicating that ATXN2 likely has a more general role as a modifier of neurotoxicity.

Our results extend the spectrum of ALS-associated proteins affected by ATXN2 to include CREST. This supports the hypothesis that therapies that target ATXN2 may be effective for a wide range of neurodegenerative diseases.

## Methods

### Yeast strains and plasmids

Yeast strains and plasmids used are listed in Tables 1 and 2. Unless otherwise stated, yeast strain BY4741 (*MAT**a** his3Δ1 leu2Δ0 met15Δ0 ura3Δ0)* was used. Yeast transformation was by the lithium acetate method [90]. L3491 bearing an integrated copy of *ADH1-HTB1-mCh* was used to check CREST nuclear localization [37]. *FLO1* (L3580) and *FLO11 (*L3582) promoter reporter strains were respectively made from DY755 and DY758. DY755 and DY758 are Ura^+^ because *FLO8,* which is disrupted in these strains, is required for regulation that inhibits the *FLO1* and *FLO11* promoters [59, 91]. We obtained pop-outs of the *HIS3* disruptions of *FLO8* by selecting for Ura^-^ revertants on FOA plates [92] and checking these for those that had become His^-^. Several such pop-outs were obtained and tested and they all behaved identically. Unless otherwise stated, all overexpression plasmids were driven by the *GAL1* promoter. Gateway technology [93] was used to make CREST entry clone p*DONR-CREST* with BP reactions between pDONR221 (Invitrogen, Cat# 12536017) and a CREST fragment amplified from p*C1-CREST* with PCR. The CREST fragment in p*DONR-CREST* was further transferred to YEp*GAL-GFP*, YCp*-GAL-GFP*, YEp*GAL-DsRED* and YIp*GAL-GFP* to respectively build YEp*GAL-CREST-GFP*, YCp*GAL-CREST-GFP*, YEp*GAL-CREST-DsRED* and YIp*GAL-CREST-GFP* with LR reactions. To increase expression of *GAL1* controlled CREST, pβ-est, containing human estrogen receptor hormone-binding domain fused to the *GAL4* DNA binding domain and the VP16 viral transcriptional activator (hER) was used.

**TABLE 1.**
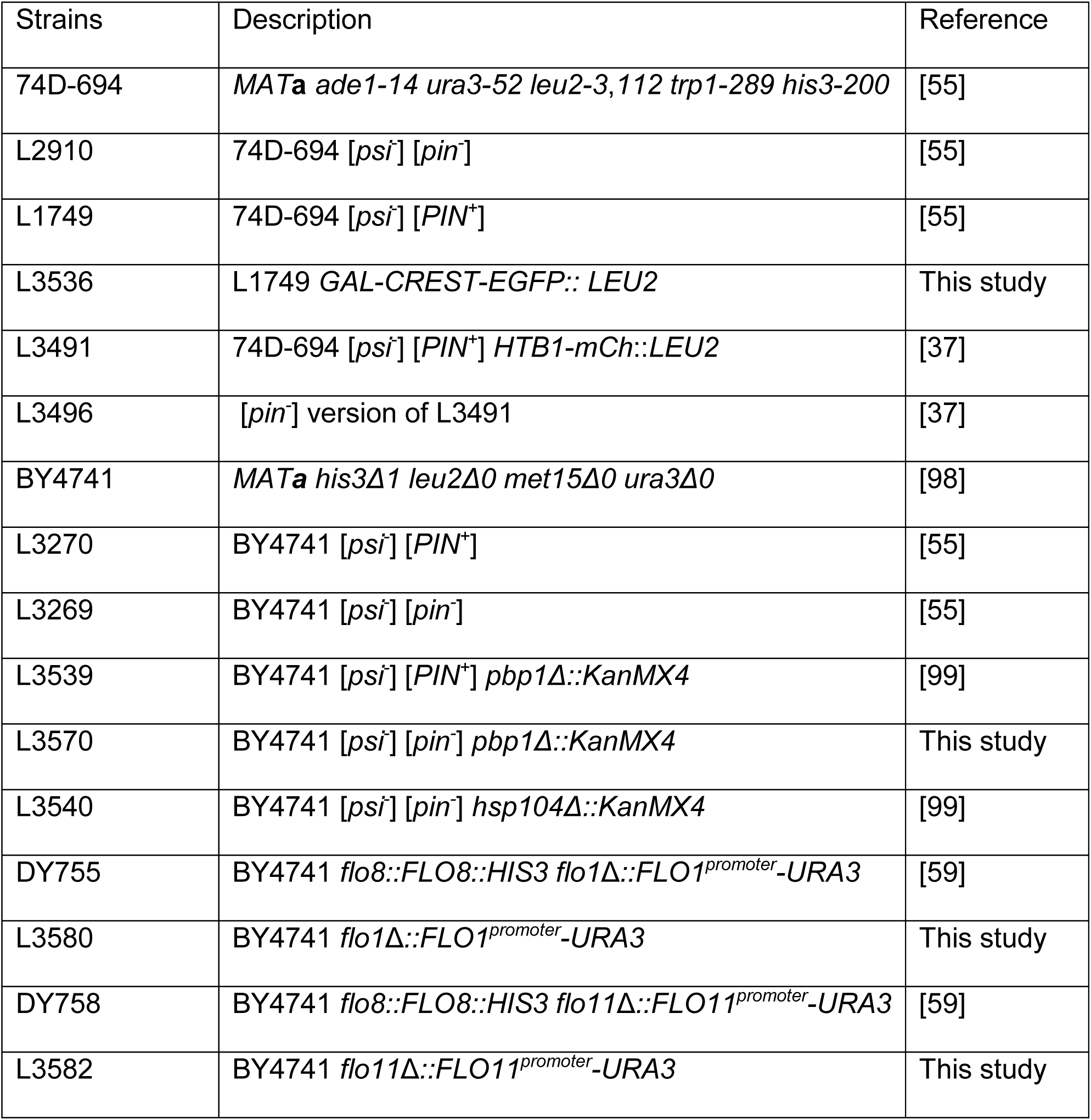
YEAST STRAINS USED.

**TABLE 2.**
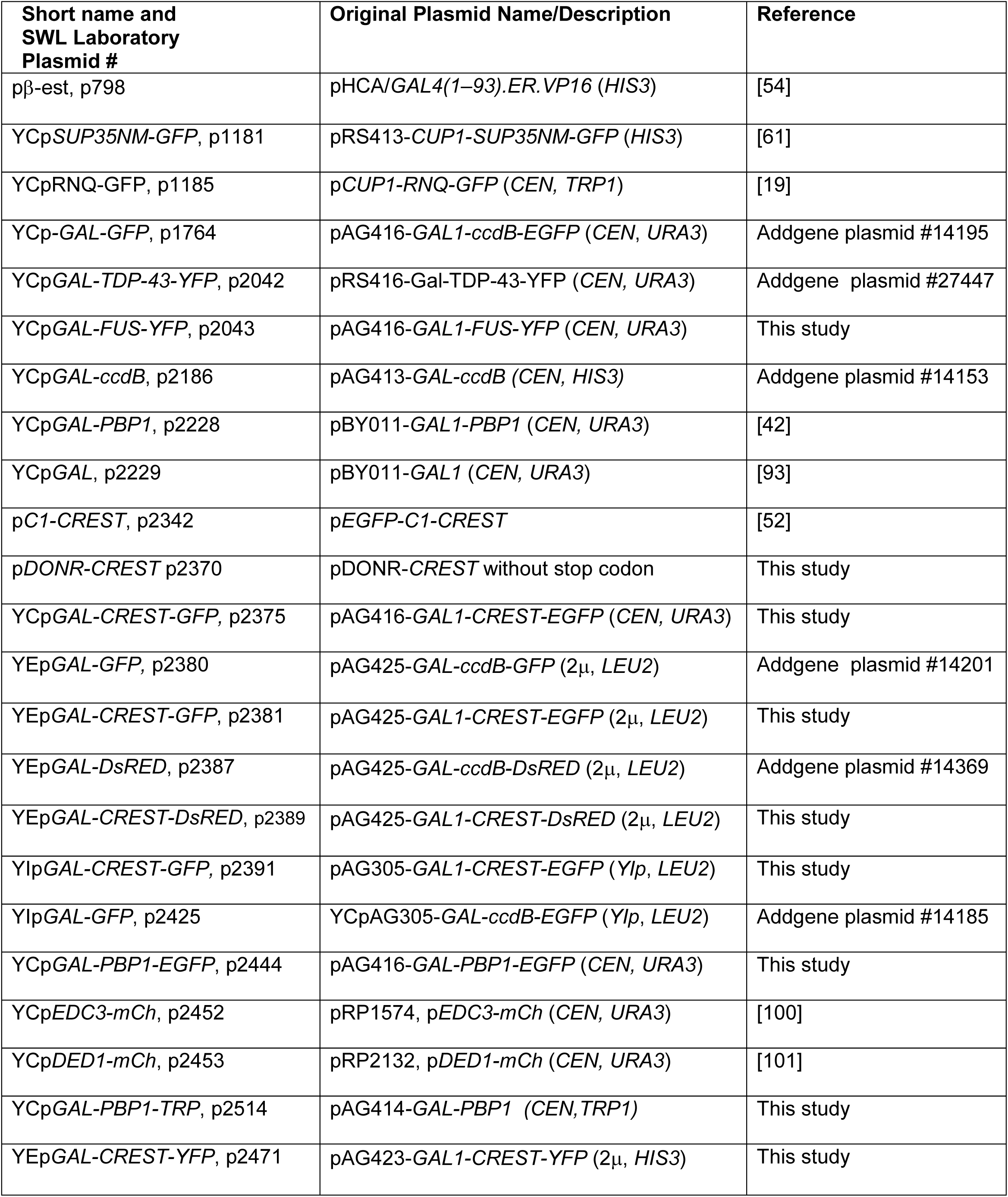
PLASMIDS USED.

### Cultivation procedures

Standard media and growth conditions were used [94, 95]. All liquid-culture experiments were carried out with mid-log-phase cells. Complex (YPD) yeast media contained 2% dextrose. Synthetic medium lacked a specific component, e.g. leucine, and contained 2% dextrose, e.g., SD-Leu (-Leu medium) or 2% galactose without (SGal-Leu) or with 2% raffinose (SRGal-Leu). Cells co-transformed with pβest, containing hER, were grown in β-estradiol (2 μM) in glucose media to turn on expression of *GAL1* controlled CREST [54],[55]. FOA medium containing 12 mg/l 5-fluoro-orotic acid and 12 mg/l uracil was used to select for Ura^-^ cells [92]. To prevent the accumulation of suppressor mutants that reverse CREST toxicity, pGAL1-CREST transformants were maintained on plasmid selective SD medium where CREST was not expressed. Patches were then replica-plated onto synthetic glycerol (2%) to identify petites that were dropped from further study. To analyze growth, non-petite transformants from SD plates were normalized in water to an OD_600_ of 2 and serially diluted 10X. Finally about 5 μl of diluted cell suspensions were spotted on plasmid selective SD, SGal or SD + β-estradiol using an MC48 (Dan-kar Corp, MA) spotter.

### Western blot analysis

CREST was detected in pre-cleared lysates separated by SDS-PAGE and blotted as described previously [96]. For analysis of proteins in total lysate vs. supernatant and pellet, the pellet from 90 μg of total lysate centrifuged at 50K rpm for 6 min was resuspended in water to the initial volume of total lysate centrifuged. The same sample volumes of total lysate, supernatant and pellet were then separated by SDS-PAGE. Blots were developed with GFP mouse antibody from Roche Applied Science (Indianapolis, IN) and PGK antibody from life technologies (Frederick, MD).

### Visualization of aggregates, nuclei and co-localization studies

Fluorescently labeled protein aggregates, were visualized in cells with a Nikon Eclipse E600 fluorescent microscope (100X oil immersion) equipped with FITC, YFP and mCh (respectively, chroma 49011, 49003 and 49008) filter cubes. To visualize nuclei, cells with the integrated HTB1-mCh nuclear marker were used [37] or nuclei were stained with 1 mg/ml 4′,6′-diamidino-2-phenylindole (DAPI) in 1XPBS for 10 min following fixing and permeabilizing cells for 1 h in 60% ethanol in PBS.

### Flocculation Assay

Flocculation was assayed by resuspending a sample of each 2 d culture grown in plasmid selective SRGal in 50 mM EDTA in water. Since flocculation requires the presence of Ca^2+^ ions [97], EDTA removed clumping allowing cells’ OD_600_ to be accurately measured. Then another sample of cells that was not treated with EDTA was appropriately normalized to OD_600_ =10. Cells were photographed after they were allowed to settle for 15 min. The level of precipitation of cells and the reduction in the OD of the supernatants (diluted 10X) is a measure of the flocculence.

### Thioflavin T Staining

Yeast cells were stained with Thioflavin T according to a protocol adapted from [31] with the addition of two extra washes in PMST [0.1 M KPO4 (pH 7.5), 1 mM MgCl2, 1 M Sorbitol, 0.1% Tween 20].

### Generation and characterization of transgenic flies

Generation of the human CREST line used in this study has been described earlier [52]. gmr-GAL4, UAS-lacZ and dAtx2 ^06490^ lines were obtained from the Bloomington Drosophila stock center. Fly stocks were raised on standard Drosophila medium at 20 °C. Crosses between the Drosophila strains were carried out at 25 °C using standard procedures. For histochemical analyses, heads of 5-day-old adult transgenic flies were dissected, collected, briefly washed in phosphate buffered saline (PBS), and fixed with 4% paraformaldehyde containing 0.1% Triton X-100 at room temperature for 2 h. After a brief wash in PBS, tissues were dehydrated by graded ethanol, cleared in butanol and embedded in paraffin. Four-micrometer thick coronal sections were stained with hematoxylin and eosin (H&E). Between 8 and 10 flies were analyzed per genotype, and 3 sections were analyzed for each fly. For Western blot analysis, heads of 5-day-old flies were lysed in Laemmli sample buffer containing 2% SDS. Anti-CREST (Proteintech, 12439-1-AP) and anti-α-tubulin (Sigma, DM1A) antibodies were used.

### Human cell culture and immunofluorescent analysis

WT CREST in pEGFP-C1 vector (Clontech) was used for CREST-GFP overexpression [52]. SH-SY5Y neuroblastoma cells were cultured in 1:1 mixture of Dulbecco’s Modified Eagle’s Medium (DMEM) and F12 medium supplemented with 10% FBS, 50 U/ml penicillin-streptomycin, and 500 μM L-glutamine (all Invitrogen). Cells were fixed with 4% cold paraformaldehyde for 15 min at RT and permeabilized with methanol. Cells were incubated with a primary anti-ATXN2 antibody (rabbit polyclonal, 21776-1-AP, Proteintech) diluted in PBS-T containing 5% goat serum at room temperature for 2 h. Fluorochrome-conjugated secondary antibody (AlexaFluor, Life Technologies) diluted 1:1,000 in PBS-T were applied at RT for 1.5 h. Nuclei were stained with DAPI and mounted on glass slides using Immunount (Thermo Scientific). Fluorescent images were taken using BX57 fluorescent microscope equipped with ORCA-Flash 4.0 camera (Hamamatsu) and cellSens Dimension software (Olympus). Figures were prepared using Photoshop CS3 software.

## Data Availability

No datasets were generated during the current study. The datasets analyzed during the current study are publicly available at https://www.yeastgenome.org/.

## Acknowledgements

We thank Ross Buchan, U. of Arizona; Bernd Bukau, U. Heidelberg; the late Susan Lindquist, MIT and Aaron Gitler, Stanford U., for plasmids and Zhiqiang Du and Liming Li, Northwestern U., for strains. In addition we thank William Eom for helpful comments on the manuscript. This work was supported National Institutes of Health Grant R01GM056350 (SWL). TAS is a Motor Neurone Disease Association Senior Non-Clinical fellow.

## Supporting Information

**S1 Fig.**
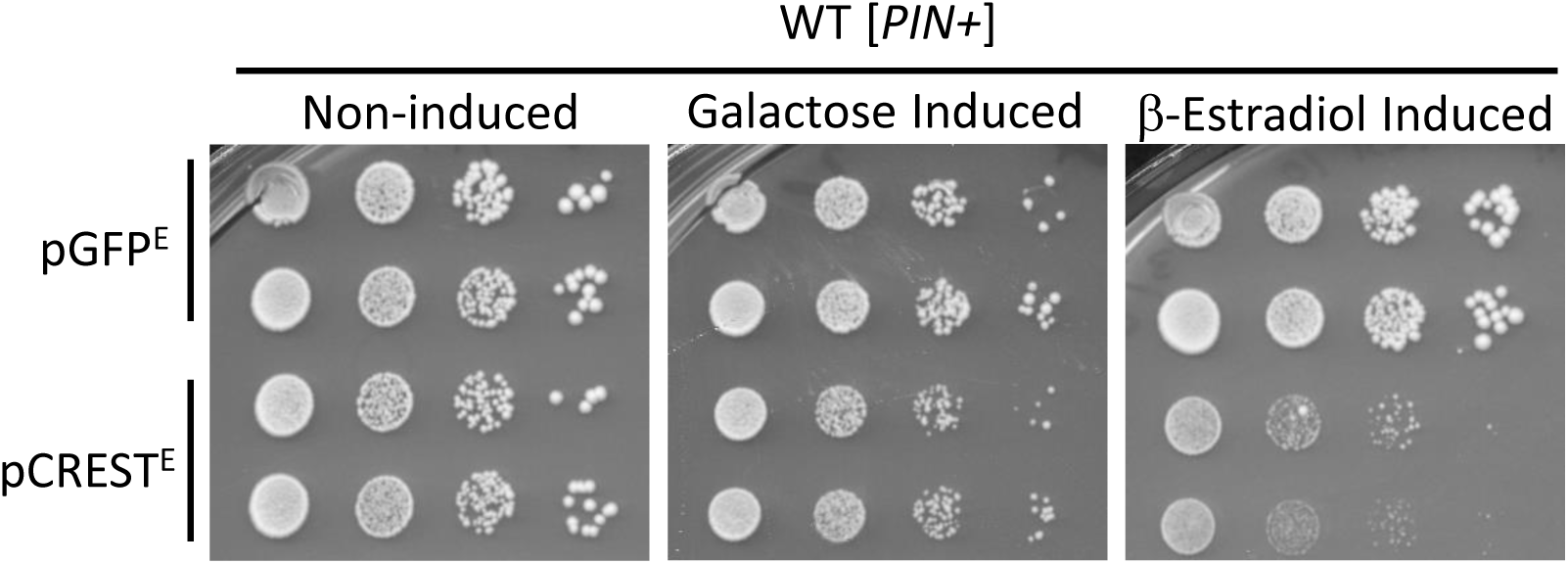
GAL-CREST-GFP is more toxic when expressed with β-estradiol than galactose. Cells transformed with pβ-est and YEp*GAL-CREST-GFP* (pCREST^E^) or the control YEp*GAL-GFP* (pGFP^E^) were serially diluted and spotted on plasmid selective glucose (Non-induced), galactose (Galactose induced) or glucose with 2 μM β-estradiol medium.

**S2 Fig.**
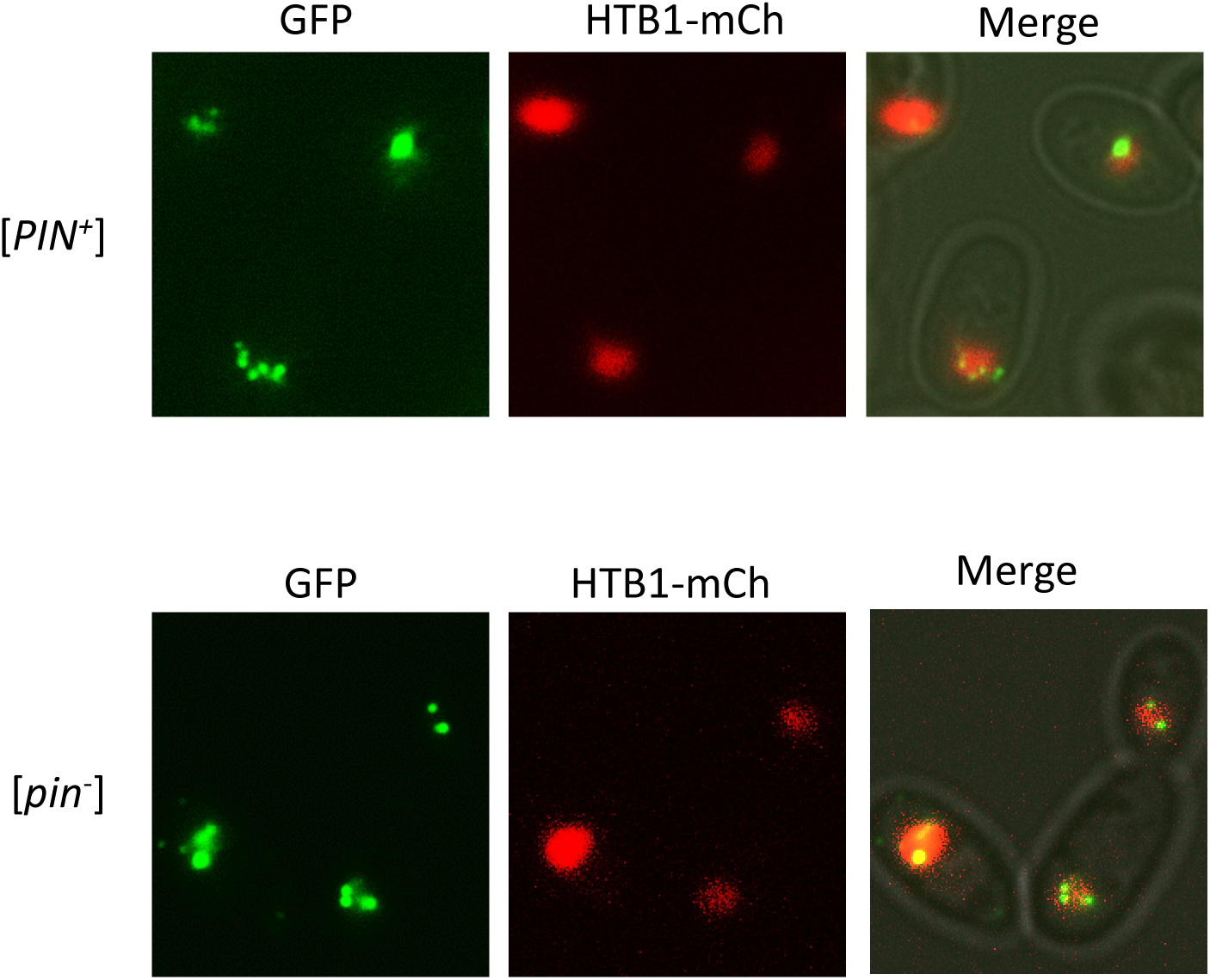
Co-localization of wild-type CREST-GFP aggregates and nuclear marker HTB1-mCh in [*PIN+*] and [*pin-*] strain. The strains L3491 and L3496 with integrated plasmid encoding ADH1-HTB1-mCh transformed with YCp*GAL*-*CREST-GFP* were grown on plasmid selective SRGal medium for 24 h and examined.

**S3 Fig.**
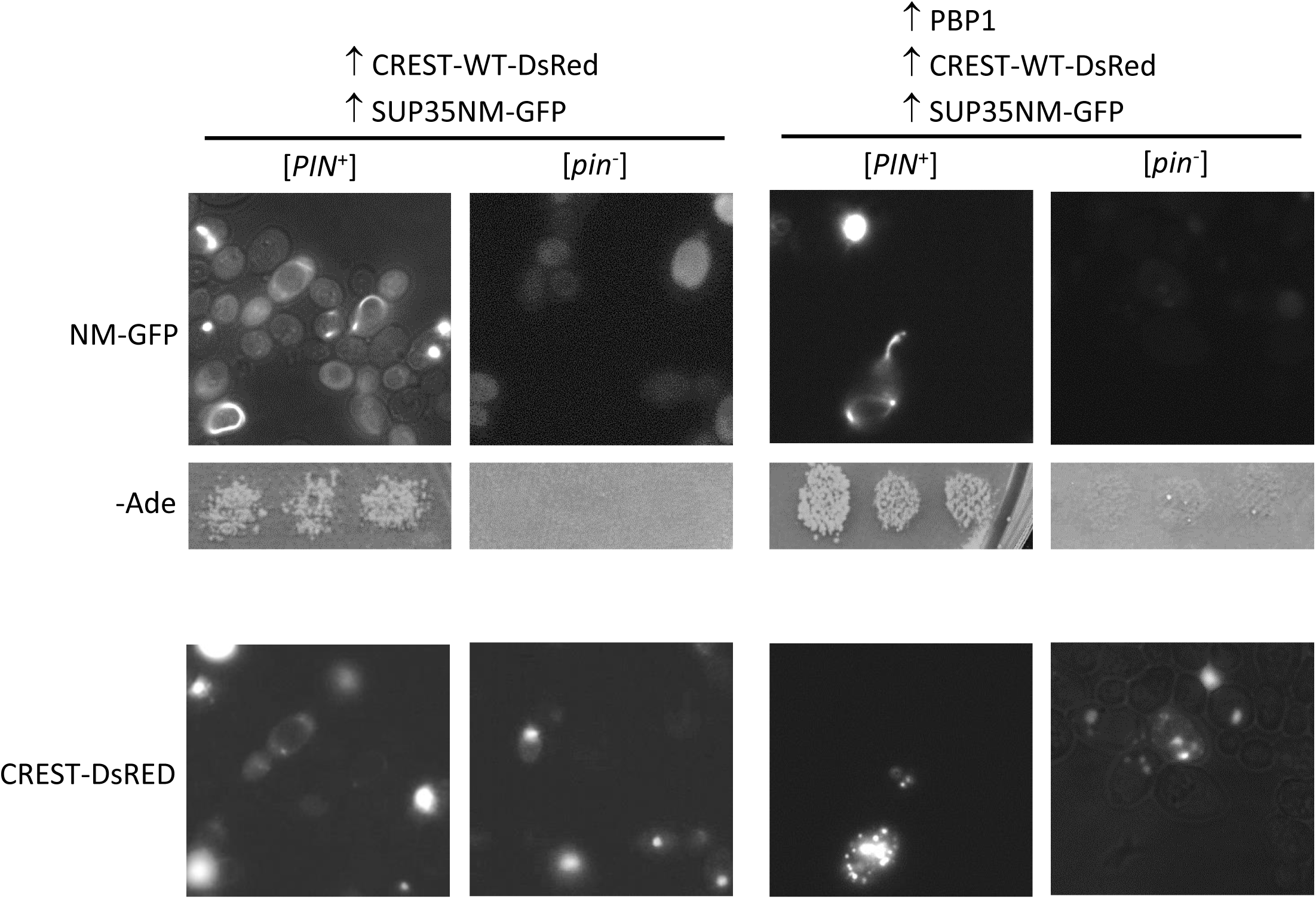
CREST does not facilitate induction of [*PSI*^+^]. Isogenic [*PIN*^+^] and [*pin*^-^] 74D-694 strains transformed with plasmids YEp*GAL-CREST-DsRED* (↑CREST-DsRED) and *CUP1-SUP35NM-GFP* (↑SUP35NM-GFP) and YCp*GAL-PBP1-EGFP* (↑PBP1) were patched on plasmid selective SGal plates with 50 μM CuSO_4_ and grown overnight. Cells were then examined under a fluorescent microscope with a GFP filter (upper) or mCherry filter (lower) and replica-plated onto medium lacking adenine (-Ade) where only [*PSI*^+^] cells can grow. Plates were photographed after incubation. Patches of three independent transformants of [*PIN*^+^] and [*pin*^-^] are shown (middle).

**S4 Fig.**
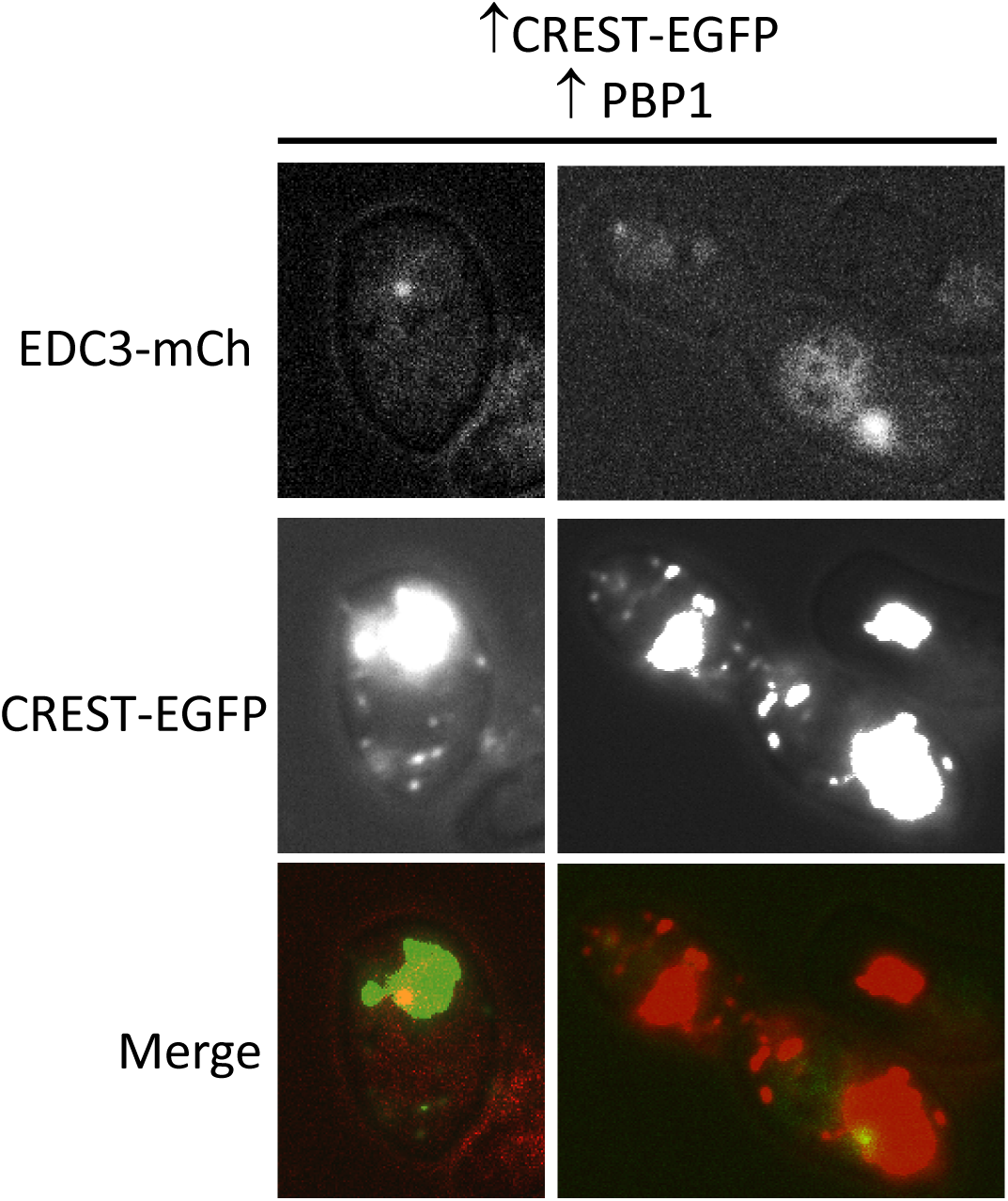
CREST-GFP foci formed by overexpression of PBP1 generally do not contain the RNP granule markers DED1 or ECD3. L1749 transformed with YEp*GAL-CREST-GFP*, YCp*GAL-PBP1-TRP* and either YCp*DED1-mCh* or YCp*EDC3-mCh* were grown on plasmid selective galactose media and examined. DED1 failed to form any dots while EDC3 dots were very rare in these stress-free cultures.

## REFERENCES

1. Bertram L, Tanzi RE (2005) The genetic epidemiology of neurodegenerative disease. J Clin Invest 115: 1449–1457.

2. Chen S, Sayana P, Zhang X, Le W (2013) Genetics of amyotrophic lateral sclerosis: an update. Mol Neurodegener 8: 28.

3. Barmada SJ, Skibinski G, Korb E, Rao EJ, Wu JY, et al. (2010) Cytoplasmic mislocalization of TDP-43 is toxic to neurons and enhanced by a mutation associated with familial amyotrophic lateral sclerosis. J Neurosci 30: 639–649.

4. Liebman SW, Chernoff YO (2012) Prions in yeast. Genetics 191: 1041–1072.

5. Wickner RB (1994) [URE3] as an altered URE2 protein: evidence for a prion analog in Saccharomyces cerevisiae. Science 264: 566–569.

6. Wickner RB, Masison DC, Edskes HK (1995) [PSI] and [URE3] as yeast prions. Yeast 11: 1671–1685.

7. Sondheimer N, Lindquist S (2000) Rnq1: an epigenetic modifier of protein function in yeast. Mol Cell 5: 163–172.

8. Derkatch IL, Bradley ME, Hong JY, Liebman SW (2001) Prions affect the appearance of other prions: the story of [PIN(+)]. Cell 106: 171–182.

9. Patel BK, Gavin-Smyth J, Liebman SW (2009) The yeast global transcriptional co-repressor protein Cyc8 can propagate as a prion. Nat Cell Biol 11: 344–349.

10. Suzuki G, Shimazu N, Tanaka M (2012) A yeast prion, Mod5, promotes acquired drug resistance and cell survival under environmental stress. Science 336: 355–359.

11. Patino MM, Liu JJ, Glover JR, Lindquist S (1996) Support for the prion hypothesis for inheritance of a phenotypic trait in yeast. Science 273: 622–626.

12. Du Z, Park KW, Yu H, Fan Q, Li L (2008) Newly identified prion linked to the chromatin-remodeling factor Swi1 in Saccharomyces cerevisiae. Nat Genet 40: 460–465.

13. Paushkin SV, Kushnirov VV, Smirnov VN, Ter-Avanesyan MD (1996) Propagation of the yeast prion-like [psi+] determinant is mediated by oligomerization of the SUP35-encoded polypeptide chain release factor. EMBO J 15: 3127–3134.

14. Nizhnikov AA, Alexandrov AI, Ryzhova TA, Mitkevich OV, Dergalev AA, et al. (2014) Proteomic screening for amyloid proteins. PLoS One 9: e116003.

15. Pezza JA, Villali J, Sindi SS, Serio TR (2014) Amyloid-associated activity contributes to the severity and toxicity of a prion phenotype. Nat Commun 5: 4384.

16. Du Z, Li L (2014) Investigating the interactions of yeast prions: [SWI+], [PSI+], and [PIN+]. Genetics 197: 685–700.

17. Derkatch IL, Bradley ME, Zhou P, Chernoff YO, Liebman SW (1997) Genetic and environmental factors affecting the de novo appearance of the [PSI+] prion in Saccharomyces cerevisiae. Genetics 147: 507–519.

18. Derkatch IL, Uptain SM, Outeiro TF, Krishnan R, Lindquist SL, et al. (2004) Effects of Q/N-rich, polyQ, and non-polyQ amyloids on the de novo formation of the [PSI+] prion in yeast and aggregation of Sup35 in vitro. Proc Natl Acad Sci U S A 101: 12934–12939.

19. Liebman SW, Bagriantsev SN, Derkatch IL (2006) Biochemical and genetic methods for characterization of [PIN+] prions in yeast. Methods 39: 23–34.

20. Derkatch IL, Liebman SW (2007) Prion-prion interactions. Prion 1: 161–169.

21. Sharma J, Liebman SW (2013) Exploring the basis of [PIN(+)] variant differences in [PSI(+)] induction. J Mol Biol 425: 3046–3059.

22. Sharma J, Liebman SW (2013) Variant-specific prion interactions: Complicating factors. Cell Logist 3: e25698.

23. Vitrenko YA, Gracheva EO, Richmond JE, Liebman SW (2007) Visualization of aggregation of the Rnq1 prion domain and cross-seeding interactions with Sup35NM. J Biol Chem 282: 1779–1787.

24. Yang Z, Hong JY, Derkatch IL, Liebman SW (2013) Heterologous gln/asn-rich proteins impede the propagation of yeast prions by altering chaperone availability. PLoS Genet 9: e1003236.

25. Serio TR (2018) [PIN+]ing down the mechanism of prion appearance. FEMS Yeast Res 18.

26. Osherovich LZ, Weissman JS (2001) Multiple Gln/Asn-rich prion domains confer susceptibility to induction of the yeast [PSI(+)] prion. Cell 106: 183–194.

27. Khan T, Kandola TS, Wu J, Venkatesan S, Ketter E, et al. (2018) Quantifying Nucleation In Vivo Reveals the Physical Basis of Prion-like Phase Behavior. Mol Cell 71: 155–168 e157.

28. Meriin AB, Zhang X, He X, Newnam GP, Chernoff YO, et al. (2002) Huntington toxicity in yeast model depends on polyglutamine aggregation mediated by a prion-like protein Rnq1. J Cell Biol 157: 997–1004.

29. Gokhale KC, Newnam GP, Sherman MY, Chernoff YO (2005) Modulation of prion-dependent polyglutamine aggregation and toxicity by chaperone proteins in the yeast model. J Biol Chem 280: 22809–22818.

30. Holmes DL, Lancaster AK, Lindquist S, Halfmann R (2013) Heritable remodeling of yeast multicellularity by an environmentally responsive prion. Cell 153: 153–165.

31. Johnson BS, McCaffery JM, Lindquist S, Gitler AD (2008) A yeast TDP-43 proteinopathy model: Exploring the molecular determinants of TDP-43 aggregation and cellular toxicity. Proc Natl Acad Sci U S A 105: 6439–6444.

32. Ju S, Tardiff DF, Han H, Divya K, Zhong Q, et al. (2011) A yeast model of FUS/TLS-dependent cytotoxicity. PLoS Biol 9: e1001052.

33. Kryndushkin D, Ihrke G, Piermartiri TC, Shewmaker F (2012) A yeast model of optineurin proteinopathy reveals a unique aggregation pattern associated with cellular toxicity. Mol Microbiol 86: 1531–1547.

34. Kryndushkin D, Shewmaker F (2011) Modeling ALS and FTLD proteinopathies in yeast: an efficient approach for studying protein aggregation and toxicity. Prion 5: 250–257.

35. Di Gregorio SE, Duennwald ML (2018) Yeast as a model to study protein misfolding in aged cells. FEMS Yeast Res 18.

36. Kurahashi H, Ishiwata M, Shibata S, Nakamura Y (2008) A regulatory role of the Rnq1 nonprion domain for prion propagation and polyglutamine aggregates. Mol Cell Biol 28: 3313–3323.

37. Park SK, Hong JY, Arslan F, Kanneganti V, Patel B, et al. (2017) Overexpression of the essential Sis1 chaperone reduces TDP-43 effects on toxicity and proteolysis. PLoS Genet 13: e1006805.

38. Treusch S, Hamamichi S, Goodman JL, Matlack KE, Chung CY, et al. (2011) Functional links between Abeta toxicity, endocytic trafficking, and Alzheimer’s disease risk factors in yeast. Science 334: 1241–1245.

39. Park SK, Ratia K, Ba M, Valencik M, Liebman SW (2016) Inhibition of Abeta42 oligomerization in yeast by a PICALM ortholog and certain FDA approved drugs. Microb Cell 3: 53–64.

40. Rosenthal SL, Wang X, Demirci FY, Barmada MM, Ganguli M, et al. (2012) Beta-amyloid toxicity modifier genes and the risk of Alzheimer’s disease. Am J Neurodegener Dis 1: 191–198.

41. Gitler AD, Chesi A, Geddie ML, Strathearn KE, Hamamichi S, et al. (2009) Alpha-synuclein is part of a diverse and highly conserved interaction network that includes PARK9 and manganese toxicity. Nat Genet 41: 308–315.

42. Elden AC, Kim HJ, Hart MP, Chen-Plotkin AS, Johnson BS, et al. (2010) Ataxin-2 intermediate-length polyglutamine expansions are associated with increased risk for ALS. Nature 466: 1069–1075.

43. Auburger G, Sen NE, Meierhofer D, Basak AN, Gitler AD (2017) Efficient Prevention of Neurodegenerative Diseases by Depletion of Starvation Response Factor Ataxin-2. Trends Neurosci 40: 507–516.

44. Figley MD, Thomas A, Gitler AD (2014) Evaluating noncoding nucleotide repeat expansions in amyotrophic lateral sclerosis. Neurobiol Aging 35: 936 e931–934.

45. Gispert S, Kurz A, Waibel S, Bauer P, Liepelt I, et al. (2012) The modulation of Amyotrophic Lateral Sclerosis risk by ataxin-2 intermediate polyglutamine expansions is a specific effect. Neurobiol Dis 45: 356–361.

46. Yu Z, Zhu Y, Chen-Plotkin AS, Clay-Falcone D, McCluskey L, et al. (2011) PolyQ repeat expansions in ATXN2 associated with ALS are CAA interrupted repeats. PLoS One 6: e17951.

47. Teyssou E, Vandenberghe N, Moigneu C, Boillee S, Couratier P, et al. (2014) Genetic analysis of SS18L1 in French amyotrophic lateral sclerosis. Neurobiol Aging 35.

48. Chesi A, Staahl BT, Jovicic A, Couthouis J, Fasolino M, et al. (2013) Exome sequencing to identify de novo mutations in sporadic ALS trios. Nat Neurosci 16: 851–855.

49. Middeljans E, Wan X, Jansen PW, Sharma V, Stunnenberg HG, et al. (2012) SS18 together with animal-specific factors defines human BAF-type SWI/SNF complexes. PLoS One 7: e33834.

50. Hodges C, Kirkland JG, Crabtree GR (2016) The Many Roles of BAF (mSWI/SNF) and PBAF Complexes in Cancer. Cold Spring Harb Perspect Med 6.

51. Staahl BT, Tang J, Wu W, Sun A, Gitler AD, et al. (2013) Kinetic analysis of npBAF to nBAF switching reveals exchange of SS18 with CREST and integration with neural developmental pathways. J Neurosci 33: 10348–10361.

52. Kukharsky MS, Quintiero A, Matsumoto T, Matsukawa K, An H, et al. (2015) Calcium-responsive transactivator (CREST) protein shares a set of structural and functional traits with other proteins associated with amyotrophic lateral sclerosis. Mol Neurodegener 10: 20.

53. Pradhan A, Liu Y (2005) A multifunctional domain of the calcium-responsive transactivator (CREST) that inhibits dendritic growth in cultured neurons. J Biol Chem 280: 24738–24743.

54. Louvion JF, Havaux-Copf B, Picard D (1993) Fusion of GAL4-VP16 to a steroid-binding domain provides a tool for gratuitous induction of galactose-responsive genes in yeast. Gene 131: 129–134.

55. Derkatch IL, Chernoff YO, Kushnirov VV, Inge-Vechtomov SG, Liebman SW (1996) Genesis and variability of [PSI] prion factors in Saccharomyces cerevisiae. Genetics 144: 1375–1386.

56. Fushimi K, Long C, Jayaram N, Chen X, Li L, et al. (2011) Expression of human FUS/TLS in yeast leads to protein aggregation and cytotoxicity, recapitulating key features of FUS proteinopathy. Protein Cell 2: 141–149.

57. Park SK, Arslan F, Kanneganti V, Barmada SJ, Purushothaman P, et al. (2018) Overexpression of a conserved HSP40 chaperone reduces toxicity of several neurodegenerative disease proteins. Prion 12: 16–22.

58. Chernoff YO, Lindquist SL, Ono B, Inge-Vechtomov SG, Liebman SW (1995) Role of the chaperone protein Hsp104 in propagation of the yeast prion-like factor [psi+]. Science 268: 880–884.

59. Du Z, Zhang Y, Li L (2015) The Yeast Prion [SWI(+)] Abolishes Multicellular Growth by Triggering Conformational Changes of Multiple Regulators Required for Flocculin Gene Expression. Cell Rep 13: 2865–2878.

60. Halme A, Bumgarner S, Styles C, Fink GR (2004) Genetic and epigenetic regulation of the FLO gene family generates cell-surface variation in yeast. Cell 116: 405–415.

61. Zhou P, Derkatch IL, Liebman SW (2001) The relationship between visible intracellular aggregates that appear after overexpression of Sup35 and the yeast prion-like elements [PSI(+)] and [PIN(+)]. Mol Microbiol 39: 37–46.

62. Satterfield TF, Jackson SM, Pallanck LJ (2002) A Drosophila homolog of the polyglutamine disease gene SCA2 is a dosage-sensitive regulator of actin filament formation. Genetics 162: 1687–1702.

63. Goncharoff DK, Du Z, Li L (2018) A brief overview of the Swi1 prion-[SWI+]. FEMS Yeast Res 18.

64. Derkatch IL, Bradley ME, Masse SV, Zadorsky SP, Polozkov GV, et al. (2000) Dependence and independence of [PSI(+)] and [PIN(+)]: a two-prion system in yeast? EMBO J 19: 1942–1952.

65. Fuentealba RA, Udan M, Bell S, Wegorzewska I, Shao J, et al. (2010) Interaction with polyglutamine aggregates reveals a Q/N-rich domain in TDP-43. J Biol Chem 285: 26304–26314.

66. Doi H, Okamura K, Bauer PO, Furukawa Y, Shimizu H, et al. (2008) RNA-binding protein TLS is a major nuclear aggregate-interacting protein in huntingtin exon 1 with expanded polyglutamine-expressing cells. J Biol Chem 283: 6489–6500.

67. Satpute-Krishnan P, Langseth SX, Serio TR (2007) Hsp104-dependent remodeling of prion complexes mediates protein-only inheritance. PLoS Biol 5: e24.

68. Seidel G, Meierhofer D, Sen NE, Guenther A, Krobitsch S, et al. (2017) Quantitative Global Proteomics of Yeast PBP1 Deletion Mutants and Their Stress Responses Identifies Glucose Metabolism, Mitochondrial, and Stress Granule Changes. J Proteome Res 16: 504–515.

69. Olbertz JH (2012) The protein SS18L1 is a potent suppressor of polyQ mediated huntingtin aggregation and toxicity. Dissertation Adviser Dr. Erich Wanker.

70. Kapeli K, Pratt GA, Vu AQ, Hutt KR, Martinez FJ, et al. (2016) Distinct and shared functions of ALS-associated proteins TDP-43, FUS and TAF15 revealed by multisystem analyses. Nat Commun 7: 12143.

71. Couthouis J, Hart MP, Shorter J, DeJesus-Hernandez M, Erion R, et al. (2011) A yeast functional screen predicts new candidate ALS disease genes. Proc Natl Acad Sci U S A 108: 20881–20890.

72. Neumann M, Sampathu DM, Kwong LK, Truax AC, Micsenyi MC, et al. (2006) Ubiquitinated TDP-43 in frontotemporal lobar degeneration and amyotrophic lateral sclerosis. Science 314: 130–133.

73. Kwiatkowski TJ, Jr., Bosco DA, Leclerc AL, Tamrazian E, Vanderburg CR, et al. (2009) Mutations in the FUS/TLS gene on chromosome 16 cause familial amyotrophic lateral sclerosis. Science 323: 1205–1208.

74. Vance C, Rogelj B, Hortobagyi T, De Vos KJ, Nishimura AL, et al. (2009) Mutations in FUS, an RNA processing protein, cause familial amyotrophic lateral sclerosis type 6. Science 323: 1208–1211.

75. Jovicic A, Paul JW, 3rd, Gitler AD (2016) Nuclear transport dysfunction: a common theme in amyotrophic lateral sclerosis and frontotemporal dementia. J Neurochem 138 Suppl 1: 134–144.

76. Archbold HC, Jackson KL, Arora A, Weskamp K, Tank EM, et al. (2018) TDP43 nuclear export and neurodegeneration in models of amyotrophic lateral sclerosis and frontotemporal dementia. Sci Rep 8: 4606.

77. Kryndushkin D, Wickner RB, Shewmaker F (2011) FUS/TLS forms cytoplasmic aggregates, inhibits cell growth and interacts with TDP-43 in a yeast model of amyotrophic lateral sclerosis. Protein Cell 2: 223–236.

78. Armakola M, Higgins MJ, Figley MD, Barmada SJ, Scarborough EA, et al. (2012) Inhibition of RNA lariat debranching enzyme suppresses TDP-43 toxicity in ALS disease models. Nat Genet 44: 1302–1309.

79. Sun Z, Diaz Z, Fang X, Hart MP, Chesi A, et al. (2011) Molecular determinants and genetic modifiers of aggregation and toxicity for the ALS disease protein FUS/TLS. PLoS Biol 9: e1000614.

80. Bakthavachalu B, Huelsmeier J, Sudhakaran IP, Hillebrand J, Singh A, et al. (2018) RNP-Granule Assembly via Ataxin-2 Disordered Domains Is Required for Long-Term Memory and Neurodegeneration. Neuron 98: 754–766 e754.

81. Li YR, King OD, Shorter J, Gitler AD (2013) Stress granules as crucibles of ALS pathogenesis. J Cell Biol 201: 361–372.

82. Maharana S, Wang J, Papadopoulos DK, Richter D, Pozniakovsky A, et al. (2018) RNA buffers the phase separation behavior of prion-like RNA binding proteins. Science 360: 918–921.

83. Li S, Zhang P, Freibaum BD, Kim NC, Kolaitis RM, et al. (2016) Genetic interaction of hnRNPA2B1 and DNAJB6 in a Drosophila model of multisystem proteinopathy. Hum Mol Genet 25: 936–950.

84. Monahan Z, Shewmaker F, Pandey UB (2016) Stress granules at the intersection of autophagy and ALS. Brain Res 1649: 189–200.

85. Becker LA, Huang B, Bieri G, Ma R, Knowles DA, et al. (2017) Therapeutic reduction of ataxin-2 extends lifespan and reduces pathology in TDP-43 mice. Nature 544: 367–371.

86. Lee KH, Zhang P, Kim HJ, Mitrea DM, Sarkar M, et al. (2016) C9orf72 Dipeptide Repeats Impair the Assembly, Dynamics, and Function of Membrane-Less Organelles. Cell 167: 774–788 e717.

87. Scoles DR, Meera P, Schneider MD, Paul S, Dansithong W, et al. (2017) Antisense oligonucleotide therapy for spinocerebellar ataxia type 2. Nature 544: 362–366.

88. Swisher KD, Parker R (2010) Localization to, and effects of Pbp1, Pbp4, Lsm12, Dhh1, and Pab1 on stress granules in Saccharomyces cerevisiae. PLoS One 5: e10006.

89. Takahara T, Maeda T (2012) Transient sequestration of TORC1 into stress granules during heat stress. Mol Cell 47: 242–252.

90. Gietz RD, Schiestl RH (2007) High-efficiency yeast transformation using the LiAc/SS carrier DNA/PEG method. Nat Protoc 2: 31–34.

91. Kobayashi O, Yoshimoto H, Sone H (1999) Analysis of the genes activated by the FLO8 gene in Saccharomyces cerevisiae. Curr Genet 36: 256–261.

92. Boeke JD, LaCroute F, Fink GR (1984) A positive selection for mutants lacking orotidine-5’-phosphate decarboxylase activity in yeast: 5-fluoro-orotic acid resistance. Mol Gen Genet 197: 345–346.

93. Alberti S, Gitler AD, Lindquist S (2007) A suite of Gateway cloning vectors for high-throughput genetic analysis in Saccharomyces cerevisiae. Yeast 24: 913–919.

94. Sherman F FG, Hicks JB (1986) Methods in Yeast Genetics; Sherman F FG, Hicks JB, editor. Plainview, New York: Cold Spring Harbor Press.

95. Sikorski RS, Hieter P (1989) A system of shuttle vectors and yeast host strains designed for efficient manipulation of DNA in Saccharomyces cerevisiae. Genetics 122: 19–27.

96. Bagriantsev SN, Gracheva EO, Richmond JE, Liebman SW (2008) Variant-specific [PSI+] infection is transmitted by Sup35 polymers within [PSI+] aggregates with heterogeneous protein composition. Mol Biol Cell 19: 2433–2443.

97. Stratford M (1989) Yeast flocculation:Calcium specificity. Yeast 5: 487–496.

98. Brachmann CB, Davies A, Cost GJ, Caputo E, Li J, et al. (1998) Designer deletion strains derived from Saccharomyces cerevisiae S288C: a useful set of strains and plasmids for PCR-mediated gene disruption and other applications. Yeast 14: 115–132.

99. Winzeler EA, Shoemaker DD, Astromoff A, Liang H, Anderson K, et al. (1999) Functional characterization of the S. cerevisiae genome by gene deletion and parallel analysis. Science 285: 901–906.

100. Buchan JR, Muhlrad D, Parker R (2008) P bodies promote stress granule assembly in Saccharomyces cerevisiae. J Cell Biol 183: 441–455.

101. Hilliker A, Gao Z, Jankowsky E, Parker R (2011) The DEAD-box protein Ded1 modulates translation by the formation and resolution of an eIF4F-mRNA complex. Mol Cell 43: 962–972.

